# Multiobjective learning and design of bacteriophage specificity

**DOI:** 10.1101/2025.05.19.654895

**Authors:** Naia Novy, Phil Huss, Sarah Evert, Philip A. Romero, Srivatsan Raman

**Affiliations:** University of Wisconsin-Madison, Biochemistry; Duke University, Biomedical Engineering; University of Wisconsin-Madison, Department of Bacteriology; University of Wisconsin-Madison, Department of Chemical and Biological Engineering

## Abstract

To better understand and design proteins, it is crucial to consider the multifunctional landscapes on which all proteins exist. Proteins are often optimized for single functions during design and engineering, without considering the countless other functionalities that may contribute to or interfere with the intended outcome. In this work, we apply deep learning to understand and design the multifunctional host-targeting landscape of the T7 bacteriophage receptor binding protein for enhanced infectivity, pre-defined specificity, and high generality in virulence toward unseen strains. We compare several different model architectures and design approaches and experimentally characterize designed phages optimized for 26 diverse tasks. We demonstrate that with multiobjective machine learning, it is possible to design complex specificities at success rates that can enable low-throughput validation of predicted hits. Our results show that the targeting capabilities of T7 are highly plastic, with opposite specificities often separated by only a few mutations. This level of tunability underscores how models trained on multifunctional data can uncover key principles of phage biology and specificity. The same modeling framework can be applied to guide the multiobjective design of other proteins or mutable biological systems, offering a general strategy for navigating multifunctional landscapes.

## Introduction

Protein function is inherently multifaceted, requiring a balance between a protein’s primary biological role while meeting demands for proper expression, folding, stability, functional regulation, and minimization of off-target interactions. Tradeoffs between these attributes constrain the sequence space accessible to evolution and engineering^1–3^. Nature, however, excels at evolving sequences that navigate these competing objectives, finely tuning biomolecules for optimal function. A striking example is the receptor-binding proteins of viruses and bacteriophages, which have evolved exquisite specificity to recognize their target hosts while avoiding non-permissive ones^4^. Molecular recognition specificity—maximizing binding toward a target substrate while minimizing interactions with anti-target substrates—demonstrates the powerful constraints shaped by evolutionary pressures. Designing proteins with multiple, often competing, functional attributes remains a significant challenge^5–8^, underscoring the need for approaches capable of learning and engineering complex phenotypes.

Bacteriophages have reemerged as promising therapeutic agents due to their ability to precisely target bacterial pathogens without the broad off-target effects associated with traditional antibiotics^9–12^. Phages achieve this targeting specificity primarily through receptor binding proteins (RBPs), which interact with molecules on the bacterial cell surface such as proteins, polysaccharides, glycolipids, and lipopolysaccharides^13,14^. Phage’s high molecular recognition specificity is a result of millions of years of evolution under strong selective pressures to recognize target motifs on bacterial surfaces in complex microbial consortia. However, unlike in natural or directed evolution, where biological systems gradually tumble toward complex fitness optima, design iterations are limited for synthetic biosystems, necessitating rapid success in designing function, specificity, and adaptability. Especially for phage, where there is a lack of reliable negative selection methods to use for directed evolution, directly designing for functional trade-offs is crucial to work toward clinical and biotechnological phage applications, requiring innovative strategies to rationally design RBPs that meet multiple, often conflicting, functional objectives.

In this work, we apply deep learning and protein language models fine-tuned with deep mutational scanning data to leverage learned understandings of the molecular determinants of bacterial recognition to design bacteriophages with tailored infectivity, specificity, and generality. Our approach utilizes a dataset comprising 26,838 variants of the T7 bacteriophage RBP, evaluated across five *Escherichia coli* strains with diverse receptor motifs. Combined, this multi-host dataset represents ∼125,000 measurements of host fitness for RBP variants ranging from 1 to 10 mutations (Supplementary Fig. 1a,b)^15,16^. Using these data, we train multi-task models that predict virulence across the five host strains from RBP sequence inputs. We implement four modeling approaches that incorporate recent advances in protein language models^17^ and structure-based deep learning^18^ (Fig. 1a) to design 7845 synthetic phages for tasks such as (1) maximizing infectivity for individual strains (5 objectives), (2) achieving specificity for arbitrary combinations of target and non-target strains (20 objectives), and (3) creating generalist phages capable of infecting all training strains (Fig. 1b). We experimentally characterized the designs and identified synthetic phage that achieve all 26 of these design objectives. The directed design of protein systems with such high complexity specificities and phenotypes is unprecedented and this work establishes a necessary framework for the simultaneous design of multiple complex protein properties.

**Fig. 1:**
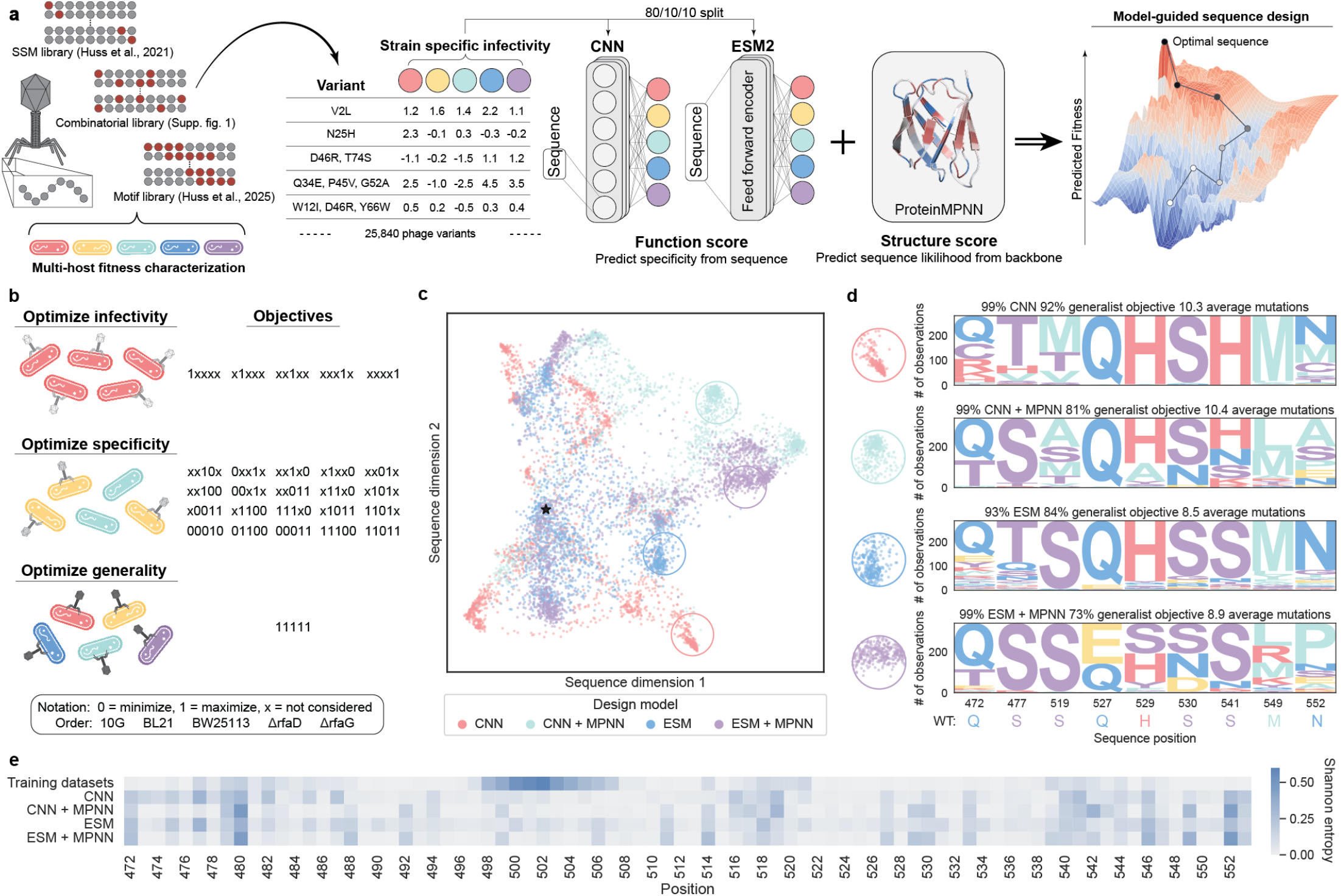
Machine learning models learn to predict and design multi-host bacteriophage function. a, Schematic of machine learning models and multi-task data used to design phages with new and enhanced functions. Each model was trained independently and paired with or without structural constraints from ProteinMPNN during sequence design. b, Phages were designed for several different objectives, broadly split up into single-task infectivity, multitask specificity, and generality. The notation describing the naming of these objectives is described here. c, Dimension reduction of the sequences designed in this study. Strong clustering is apparent by the modeling approach used to design sequences. The wild-type sequence is indicated with a star. As one example, clusters of sequences that commonly belong to each model are circled and divergent regions of the sequence motifs for these highlighted sequences are shown in d. e, Diversity of mutation sampling at each position for designs from each model and the original training datasets, measured by Shannon Entropy of mutation frequencies. Mean entropies for each subset are 0.079, 0.079, 0.067, 0.092, 0.075, following the order shown.

## Results

### Machine learning-directed design of bacteriophages

A phage’s RBP dictates its species targeting and strain selectivity, but the molecular details of these interactions are largely unknown. In our previous work, we generated large-scale mutagenesis data mapping variants in the terminal 82 residues of the T7 RBP across five *E. coli* strains^15,16^. These hosts were selected due to their known differences in susceptibility to wild-type T7. Data from this work represent both single mutations in the RBP^15^ and higher-order mutations selected with guidance from metagenomic databases^16^. To augment the information density of the training data, we created an additional dataset for multi-host fitness of bi- and tri-mutant RBPs with combinations of confirmed single mutants that drove changes in the host range (Supplementary Fig. 1a,b). These combined data provide a valuable resource for understanding how the T7 RBP sequence influences infectivity and selectivity.

We apply supervised sequence-function models^19,20^ to learn the sequence-specificity mapping from T7 RBP variants to strain-specific virulence. To evaluate which modeling approaches were best suited for these datasets, we examined the extrapolation performance of different approaches by training models on T7 RBP sequences with 1-3 mutations and measuring their predictive performance on data for held-out higher-order mutants with an average of ∼6 mutations. We explored three model architectures: a simple linear regression (LR) model, a convolutional neural network (CNN) that learns non-linear features across an input sequence^20,21^, and a transformer-based protein language model (ESM2)^17^ fine-tuned for phage fitness prediction, all trained using a standard mean square error (MSE) regression loss function^22^. We also compared a custom loss function, termed a thresholded MSE loss, which accounts for inaccurate fitness measurements caused by the dropout of non-functional variants during selection (Supplementary Fig. 2a). We found that more complex modeling approaches, such as neural networks (CNN and ESM2) and custom loss functions, improved the model’s predictive accuracy over linear regression or standard MSE regression (Supplementary Fig. 2b). Based on these results, we chose to move forward with the CNN and fine-tuned ESM2 architectures in combination with the thresholded MSE loss function.

To guide the design of synthetic phages, we compiled a single dataset consisting of all data (train and extrapolation sets from above) and trained ensembles of either CNN or ESM2 models with independent random parameter initializations and training and validation datasets, using 100 or 10 models in the CNN and ESM2 ensembles, respectively. The final CNN and ESM2 ensembles showed high accuracies (weighted Spearman ∼0.8) when evaluated on a shared held-out test set of 10% of all data, which included sequences not seen by any models during training and across all three training datasets (Supplementary Fig. 3a-c). We used these models to guide the phage design process by employing simulated annealing to search for optimal RBP sequence configurations^20,23,24^ (Fig. 1a). We also tested if incorporating structural information from ProteinMPNN could boost performance by increasing the likelihood that a given design will fold into the wild-type backbone structure^18^. Altogether, we designed synthetic phages using four approaches: CNN, CNN + MPNN, ESM, ESM + MPNN.

Employing each of these four sequence generation methods, we optimized phages for 26 different design objectives: (1) high infectivity for each of the five *E. coli* strains individually, (2) 20 different specificity combinations, and (3) generalist phages that infect all strains (Fig. 1b). These phage variants contained 3, 6, 9, and 12 mutations compared to the wild-type T7 RBP which enabled assessment of model performances at diverse mutational distances. Altogether, we designed, synthesized, and tested a total of 7845 designed phages. Each design approach led to distinct mutational patterns in the resulting sequences, despite being trained on the same experimental data (Fig. 1c,d, Supplementary Fig. 4a). We highlight the mutational preferences between models by showing the sequence motifs for divergent positions from sequences subsampled from distal sequence clusters with a high occurrence of sequences from each model (Fig. 1d). Notably, the addition of structural constraints from ProteinMPNN reduced an effect observed in the structure-agnostic CNN and ESM models in which low levels of background mutations were observed throughout the protein (Fig. 1e). Inclusion of structural constraints also increased the preponderance of conservative mutations, measured by mean BLOSUM80 score^25^ (Supplementary Fig. 4b, p<5e-60). The bias of ProteinMPNN toward conservative mutations could restrict the portion of sequence space accessible to design for complex specificities.

One concern when using machine learning to generate new protein variants is that models will regenerate sequences preexisting in the training data. Although each model generally favored mutations at similar positions in the T7 RBP, the overall distributions of mutations for the ML-designed phages were highly divergent from the training data (Fig. 1e). Even though we did not directly prohibit generated sequences from overlapping with the training data, only one designed bacteriophage was an exact match for a previously observed phage. We found a small relationship between the mutational distance and frequency of conservative mutations, but this effect size was minimal other than comparing three mutants to higher-order mutants (Supplementary Fig. 4c, p<5e-30). This suggests that after including a few choice mutations, additional mutations are on average slightly more conservative.

We constructed a pooled library of synthetic phages using ORACLE^15^. ORACLE is a phage genome engineering tool we previously developed that uses Cre recombinase-guided insertion of a variant library at a targeted genomic locus (Fig. 2a). The resulting phage population represents the pre-selection library of RBP variants, which was then incubated independently with each strain of interest in triplicate. We sequenced pre- and post-selection libraries and calculated the relative change in abundance of each library member to determine phage fitness on each host. These calculations yield continuous measurements of relative virulence for each phage in the library against each host of interest.

**Fig. 2:**
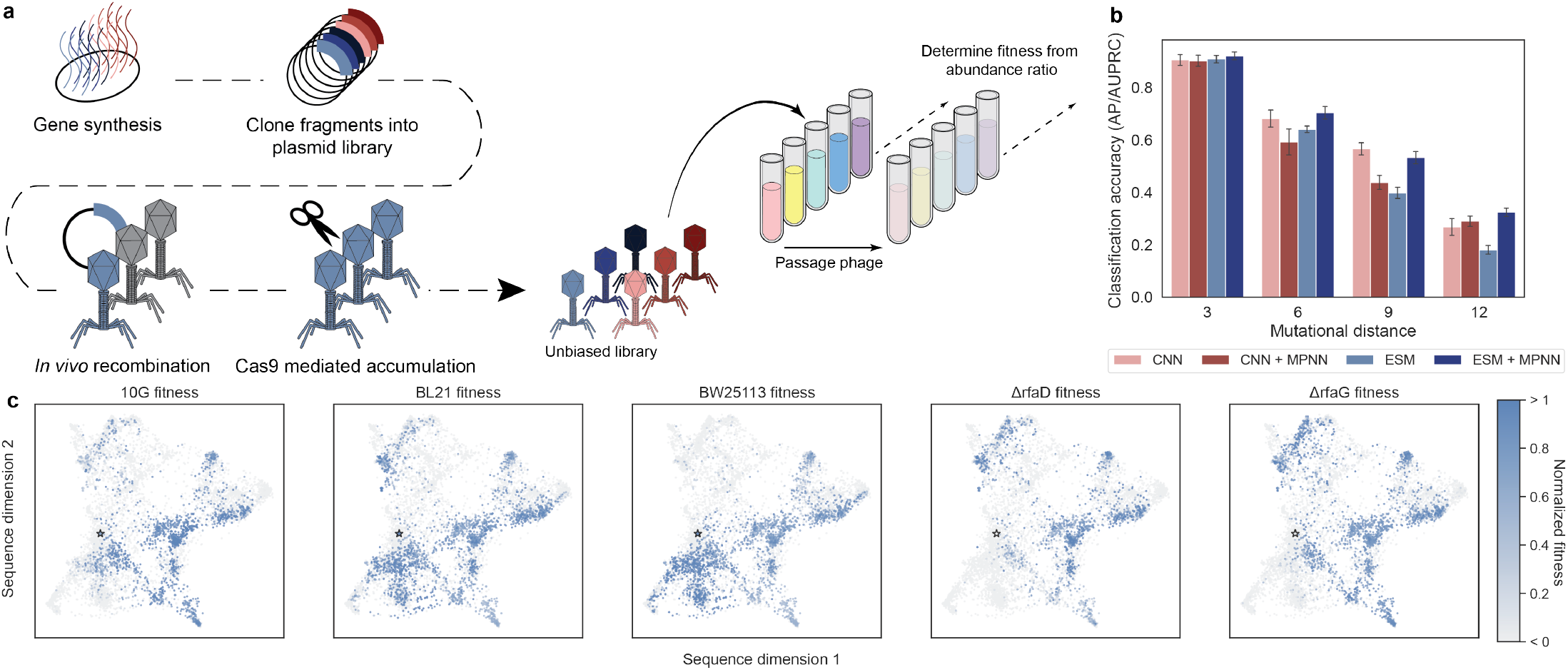
Generation and multi-host characterization of ML-designed phages. a, Schematic of bacteriophage cloning and the selection and sequencing assay. b, Overall classification accuracy (AUPRC/AP) for predicting if a designed phage is successful for its target objective. Success is defined as activity above the limit of detection (LOD) for infectivity, higher virulence on all target hosts than anti-target hosts for specificity, and the ability to infect all strains screened for generalists. AP is calculated for each objective and the mean is shown, with error bars representing the standard error over all objectives. c, Designed phages show variant-specific preference towards different hosts when visualized with dimension reduction. Fitness scores are normalized between 0 and 1, where less than 0 represents non-functional phages below LOD and greater than 1 indicates phages are above the 98th percentile for the relevant host. Wild-type is indicated by a star on each subplot.

We evaluated the predictive capacity of models using the average precision (AP/AUPRC) classification^26^ metric based on a model’s ability to classify whether phages successfully met their given objective, defined as the ability to infect the target host for infectivity, higher virulence on target hosts than anti-target (minimized) hosts for specificity, and ability of phage to infect all screened hosts for generalist designs. All models demonstrated high classification accuracy at low mutational distances, and ESM + MPNN had the highest accuracy at all mutational distances (Fig. 2b). To assess where there were differences in the mutations for non-functional phage compared to all designed phage, we compare the distributions of mutation frequencies between these two groups using Kullback-Leibler Divergence (KLD)^27^. We found that each model made mistakes at different positions, due to increased KLD at different regions in the primary sequence (Supplementary Fig. 5a). This contrasts with the effects of mutational distance, for which mistakes were generally made in similar positions regardless of the number of mutations (Supplementary Fig. 5b). In both cases, the location of mutations that were predicted to be functional but were actually deleterious were primarily made at positions where the model less frequently introduced mutations (Supplementary Fig. 5a,b)

We mapped the measured fitness values onto a dimension reduction of the designed sequences to visualize the fitness relationships between designed bacteriophages (Fig. 2c). Through this analysis, distinct patterns are apparent in the fitness and specificity landscapes, such as clusters of sequences with multi-host specificity and high similarity in the sequence-fitness effects for susceptible strains (10G, BL21, BW25113) or resistant strains (BW25113*ΔrfaD*, BW25113*ΔrfaG*). This qualitative analysis showcases the malleability of phage fitness and the ability to design phages with diverse specificity profiles.

### Machine learning enables the design of diverse phages with high virulence

When utilized in therapeutic or biosanitation applications, bacteriophages are often administered as cocktails of diverse phages, which helps restrict target pathogen evolution^11,12,28,29^. It is important for each phage in these cocktails to show high virulence toward the target host. Thus, our first design objective, which we denote as “infectivity”, was to design phages with high virulence on individual target *E. coli* strains (10G, BL21, BW25113, ΔrfaD, and ΔrfaG).

We define success in designing phage infectivity as the ability of a phage to infect its intended host above the limit of detection. The majority (80-95%) of designed triple mutants successfully infected their target strain, while designs at high mutational distances showed reduced success rates (Fig. 3a). Nonetheless, over 30% of the 12-mutant designs generated by the CNN and ESM+MPNN models retained the ability to infect the target strain. When ignoring evolutionary constraints, sequences designed with the CNN were more likely to be functional than those designed with ESM at an edit distance of 9 (p<0.01), but not other distances (Supplementary Fig. 6a). Focusing instead on the impact of structural constraints, there was no significant difference in infectivity design outcomes at any mutational distance between models that included ProteinMPNN and those that did not (Supplementary Fig. 6b). No improvement in design success when including ESM or MPNN may be a result of overemphasizing protein stability over dynamic function^30^ or could indicate that the sequence-function models had sufficient data to model single-host virulence and adding additional constraints hindered success.

**Fig. 3:**
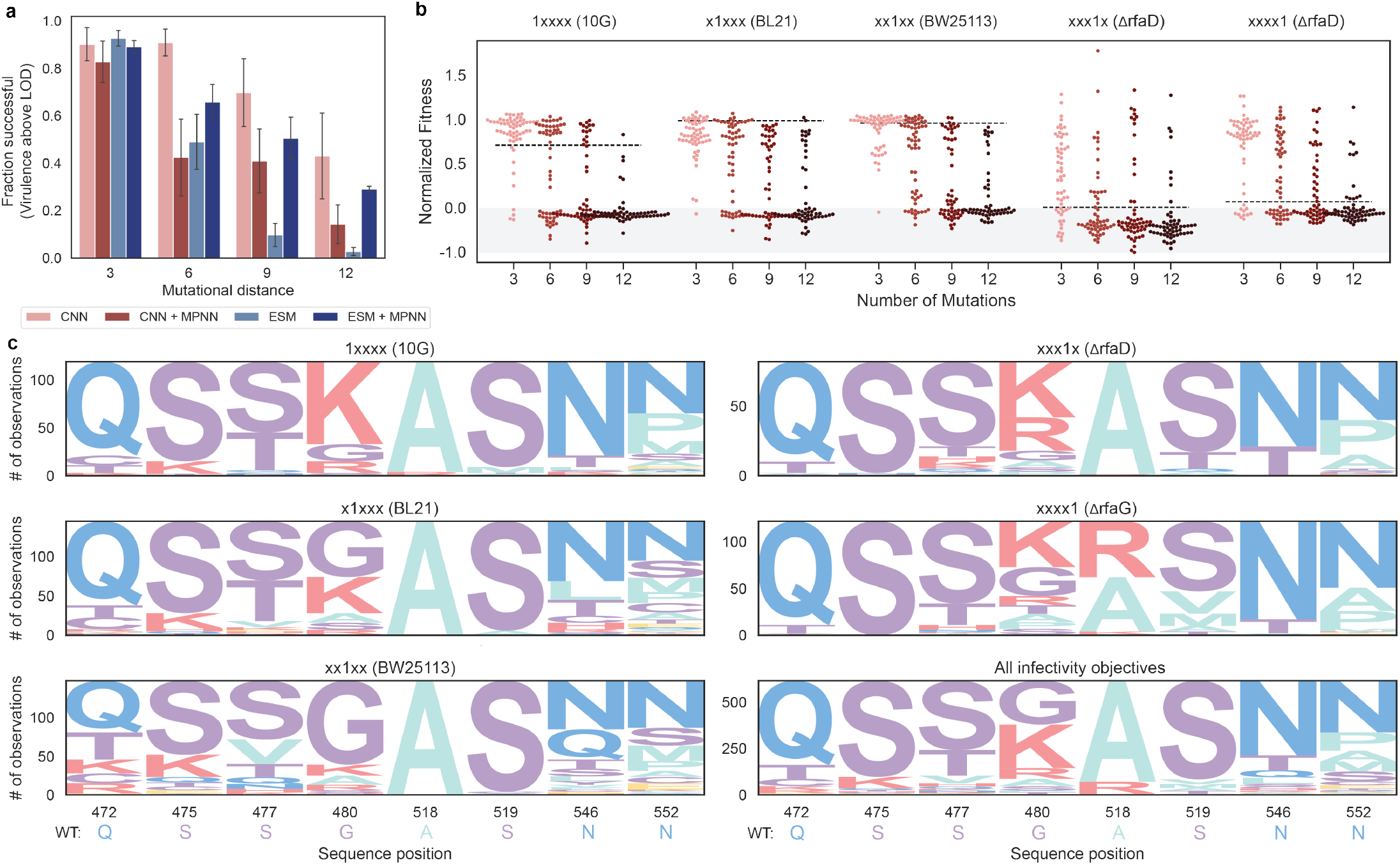
ML-directed optimization enables efficient design of phage virulence. a, Success rate for designing infectivity by model and mutational distance, where success is defined as virulence above the LOD for each bacterial host. Standard error bars are calculated using the fractional success for all five infectivity objectives. b, Normalized fitness for phage designed for each infectivity objective (see methods for normalization approach). Points are colored according to the mutational distance, also labeled on the x-axis. The black dashed line is the wild-type score for each host strain. Points in the grey region represent non-functional phages. c, Sequence logos for a subset of divergent positional frequencies for each infectivity objective.

For hosts with high susceptibility to wild-type T7 (10G, BL21, BW25113), designed phages show minimal improvement compared to wild-type T7 (Fig. 3b). This is likely because T7 is already highly optimized for these hosts, leaving limited room for further enhancement. In contrast, wild-type phage is greater than four orders of magnitude less infective on some resistant hosts (ΔrfaD and ΔrfaG) compared to susceptible hosts^15^. Regardless of this natural deficiency, many phages designed for these resistant hosts are several orders of magnitude more virulent than wild-type T7. In general, there are minimal mutational motifs for the successful variants within each objective (Fig. 3c). We found that only a few mutations are necessary to improve phage and increasing the mutational distance was unlikely to enhance virulence further (Fig. 3b, Supplementary Fig. 6c). Considering that we observe high success without the emergence of core motifs, it appears that there are many potential optima to modify the T7 RBP for enhanced virulence on single hosts.

### Directed design of high-order bacteriophage specificity

Designing phages with high-order targeting specificities is challenging and not easily achieved through traditional directed evolution methods that use alternating rounds of positive and negative selection^8,31^. Multi-task machine learning models, which map amino acid sequences to multiple properties, offer a promising approach for designing proteins with precisely defined combinations of properties. We tested our models by designing phages for 20 different specificity objectives, each targeting a unique combination of 2-5 target and anti-target strains, for which predicted virulence was maximized or minimized, respectively (Fig. 1b). These objectives were selected by computationally designing phages for all possible permutations of 2-5 part specificities and selecting a subset of diverse objectives with designs predicted to have the intended specificity. Design success for this specificity task is defined by a phage showing higher relative virulence on all target strains than all anti-target strains (scores are calculated from selection and sequencing assay). We found that the ESM-based models outperformed their CNN counterparts at edit distances of 6 and 9 (Supplementary Fig. 7a, p<0.01). Inclusion of structural constraints with ProteinMPNN improved design success rates for specificity by 9, 3, and 11% at mutational distances of 6, 9, and 12 (Supplementary Fig. 7b, p<0.002). Overall, the best model for designing specificity was ESM+MPNN, which had higher design success compared to all models at 9 mutations and CNN and ESM base models at 12 mutations (Fig. 4a, p<0.02).

**Fig. 4:**
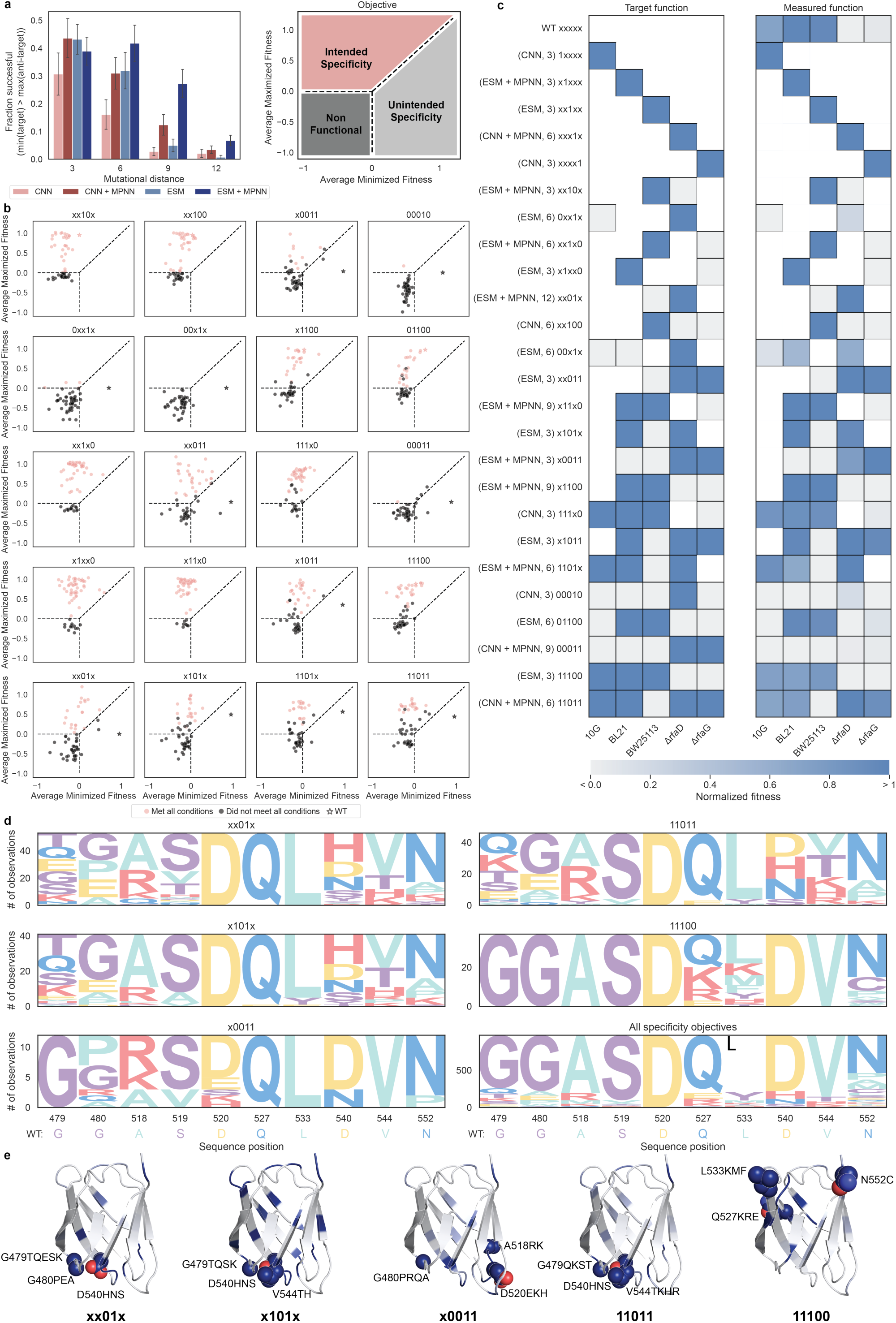
Directed design of bacteriophage targeting specificity. Refer to Fig. 1b for the legend of specificity codes. a, Fraction of phages successfully designed for specificity by model and mutational distance. Where specificity design success is defined as a phage showing higher virulence on all targets than all anti-target hosts. Standard error bars are calculated using the fractions for all 20 specificity objectives. b, Average fitness of three mutant phages on the maximized and minimized strains for each specificity objective. Red points meet all criteria for a particular specificity and black points were unsuccessful for one or more conditions. Wild-type fitness is indicated by a star. c, Top performing phages for each of the design objectives. The left subplot is the intended function and the right is the normalized fitness (see methods) as measured in the pooled screen. Spaces that are colored white were not considered in the design task. d, Sequence logos (including wild-type) for a subset of divergent positional frequencies for selected specificity objectives. These residue positions were selected due to their differences in the selected set of objectives. e, Structures of T7 receptor binding domain colored by mutational frequencies at each position for successful designs from selected specificities. The wild-type residues for the three most mutated positions of each objective are shown as spheres and mutations that were present in >8% of successful designs are annotated.

We denote each specificity objective as a sequence of five characters representing the five strains: 10G, BL21, BW25113, ΔrfaD, and ΔrfaG. In this notation, a “1” indicates that virulence toward that strain was maximized, a “0” indicates that virulence was minimized, and strain positions marked as “x” were not considered (Fig. 1b). For example, the task “x101x” specifies an objective that maximizes fitness for BL21 and ΔrfaD, minimizes fitness for BW25113, and does not consider 10G or ΔrfaG. For a more granular assessment of the specificity of designed phages for each objective, we plotted the average fitness of the target strains against the average fitness of the anti-target strains. In this analysis, successful specificity designs are expected to occupy the upper left sector, and the complexity of the objectives increases from left to right across the subplots. Many of these designed phages demonstrated the intended specificity profile, with some showing high virulence on all target strains and infection below the limit of detection for anti-target strains (Fig. 4b, Supplementary Fig. 8a-c).

Based on results from our high-throughput phage selection assay, we successfully designed phages for all 20/20 specificity objectives. For 19 of these objectives, our designs achieved diverse specificities with only 3 mutations from wild-type T7, indicating high adaptability in the natural T7 RBP (Fig. 4b). The 00×1x objective only had successful designs at higher mutational distances (Supplementary Fig. 8a). In several instances (x0011, 00010, 0xx1x, 00×1x, xx011, 00011, x1011, x101, 1101x, 11011) our designs achieved the complete opposite specificity as wild-type T7. Overall, it was very uncommon for designs to have a specificity opposite to the intended task, and the more common mode of failure was for a designed phage to be completely non-functional, which is identified by variant dropout in the post-selection phage population.

We observed that some specificity objectives were much more challenging to design (00010, 0xx1x, 00×1x, 00011), and most of the designed phages for these objectives were simply nonfunctional. This is in part because, during optimization for these tasks, many designed sequences were predicted to have a large gap in scores between maximized and minimized objectives, but overall low scores for both the target and anti-target strains. Over-minimization toward anti-target hosts is likely also a contributing factor to the progressive decline in success rates for higher complexity objectives (Supplementary Fig. 9). This over-minimization of negative function can be remediated through the inclusion of evolutionary or structural constraints (Supplementary Fig. 7a,b). Retrospectively, this effect could have also been mitigated by adjusting the scoring function to co-optimize for high fitness on target strains in addition to a large gap between maximized and minimized strains. Because the most common failure mode when designing specificity was a complete loss of virulence, altering the optimization function or including structural constraints could enhance design success in other specificity design projects.

We calculated the target and measured fitness for the best-designed phages across all models and edit distances by normalizing the fitness scores for each strain between the lower limit of detection and the 98^th^ percentile observation (see methods). Scores for strains that were not considered during design are concealed for clarity. Each design model produced the best mutant phage for at least one specificity objective and the best mutants were found at each mutational distance for at least one objective (Fig 4c). While we do observe phages that are selective for each task, objectives 0xx1x, 00×1x, 00010, and 00011 had the weakest specificities. These all share the common trend of simultaneously maximizing resistant strains and minimizing susceptible strains, which is an inherently challenging design task.

To investigate the mutational drivers underlying diverse phage specificities, we analyzed a group of design objectives that share a common motif: minimizing BW25113 while maximizing ΔrfaD. This set includes x0011, xx01x, x101x, and 11011. Among these, xx01x, x101x, and 11011 consistently favor charged or polar substitutions at positions 479, 540, and 544—three of the most frequently mutated sites in phages optimized for these objectives (Fig. 4d,e). In contrast, x0011 tends to preserve wild-type residues at these positions, instead favoring mutations at positions 480, 518, and 520. Since x0011 differs from the others principally in its additional requirement to minimize BL21 virulence, these patterns suggest the existence of at least two distinct optima within the core motif: one that tolerates or enhances activity against BL21, and one that actively suppresses it. More broadly, the emergence of divergent mutational preferences among objectives sharing the same BW25113/rfaD motif highlights the presence of multiple local optima and illustrates the ability of multi-task sequence-fitness models to navigate these trade-offs when engineering phage specificity across hosts.

Structurally, some objectives show a higher frequency of mutations at the exterior loops of the RBP, which is expected to be the primary receptor binding interface receptor^15,32,33^. However, many mutations were observed across the entire RBP domain for all objectives (Supplementary Fig. 10a-e). Of note, for objective 11100, the top three most frequently mutated positions were located on the interior loops, which are distal the exterior loops that are expected to primarily interact with the cell surface (Fig. 4e, Supplementary Fig. 10e). The presence of key mutations in non-intuitive places in the protein structure highlights the importance of mutational scanning beyond key domains.

### Machine learning-directed design of generalist phages capable of infecting new hosts

A major constraint of natural phages is their inherently narrow host range, which significantly reduces their utility against non-predisposed or rapidly evolving bacterial populations. This limitation is particularly problematic in clinical settings, where timely and precise strain identification is often challenging, resulting in the frequent use of broad-spectrum antibiotics. Rapidly engineering generalist bacteriophage with a broad host range is therefore of high interest. Ideally, a researcher could start with a model phage system and identify mutations that render novel or enhanced virulence toward new targets. In line with this goal, the final design objective was generality.

These phages were optimized to exhibit high fitness on each of the five strains for which we had data. Characterization of synthetic generalists revealed that these phages were frequently capable of infecting all training strains with approximately 80% success rates at an edit distance of 3 (Supplementary Fig. 11a,c). At edit distances of 6 and 9 mutations, the CNN and ESM + MPNN design approaches consistently produced generalist phages more frequently than the other two approaches (p≤0.025), each achieving >50% success for infecting the five target strains at 9 mutations (Supplementary Fig. 11c).

While high success rates were observed in designing generalists towards the training strains, the ideal advantage of designing generalist function lies in the potential for phages optimized toward known strains to infect a broader range of strains with unknown susceptibility profiles, accelerating screening for novel fitness. To evaluate the true promiscuity of these phages, we challenged the ML-designed phages against eight additional *E. coli* strains with alternative modifications to the surface receptors, including deletions to genes involved in lipid polysaccharide (LPS) modification (ΔrfaC, ΔrfaF, ΔrfaJ, ΔrfaI, ΔlpcA), iron transporter secondary receptors (ΔtonA and ΔtonB), and full deletion of the LPS primary receptor (ClearColi).

The receptor deletion strains are expected to have variable degrees of truncations to the LPS, with ΔrfaC, ΔlpcA, and ΔrfaD resulting in expected truncations directly above the KDO sugar, followed by ΔrfaF, ΔrfaG, ΔrfaI, and ΔrfaJ each truncating one additional level of sugars above this KDO base relative to each other^16,34^. ΔtonA and ΔtonB should show similar variant-host susceptibility as BW25113 as these deletions have the same primary LPS receptor as BW25113^35^. Many generalist phages exhibited high virulence on these non-training strains even without being directly designed for these tasks (Supplementary Fig. 11b,c). Approximately 35% of phages designed by the CNN or ESM + MPNN models were functional on all 13 *E. coli* receptor deletion strains at an edit distance of 9 (Fig. 5a). Phages that did not infect all 13 strains were typically only unable to infect 1-2 non-training strains at low mutational distances (Fig. 5b). At higher mutational distances, the most common failure mode was an inability to infect any of the 13 strains, suggesting a complete loss of phage viability (Supplementary Fig. 11d). The frequency of occurrence of generalist behavior among all phages was ∼2x higher for phages directly optimized as generalists than phages designed for infectivity, and ∼30x higher than specialist phages (Fig. 5c). This significant improvement in promiscuity emphasizes the advantage of machine learning-directed generalist design to optimize for new tasks.

**Fig. 5:**
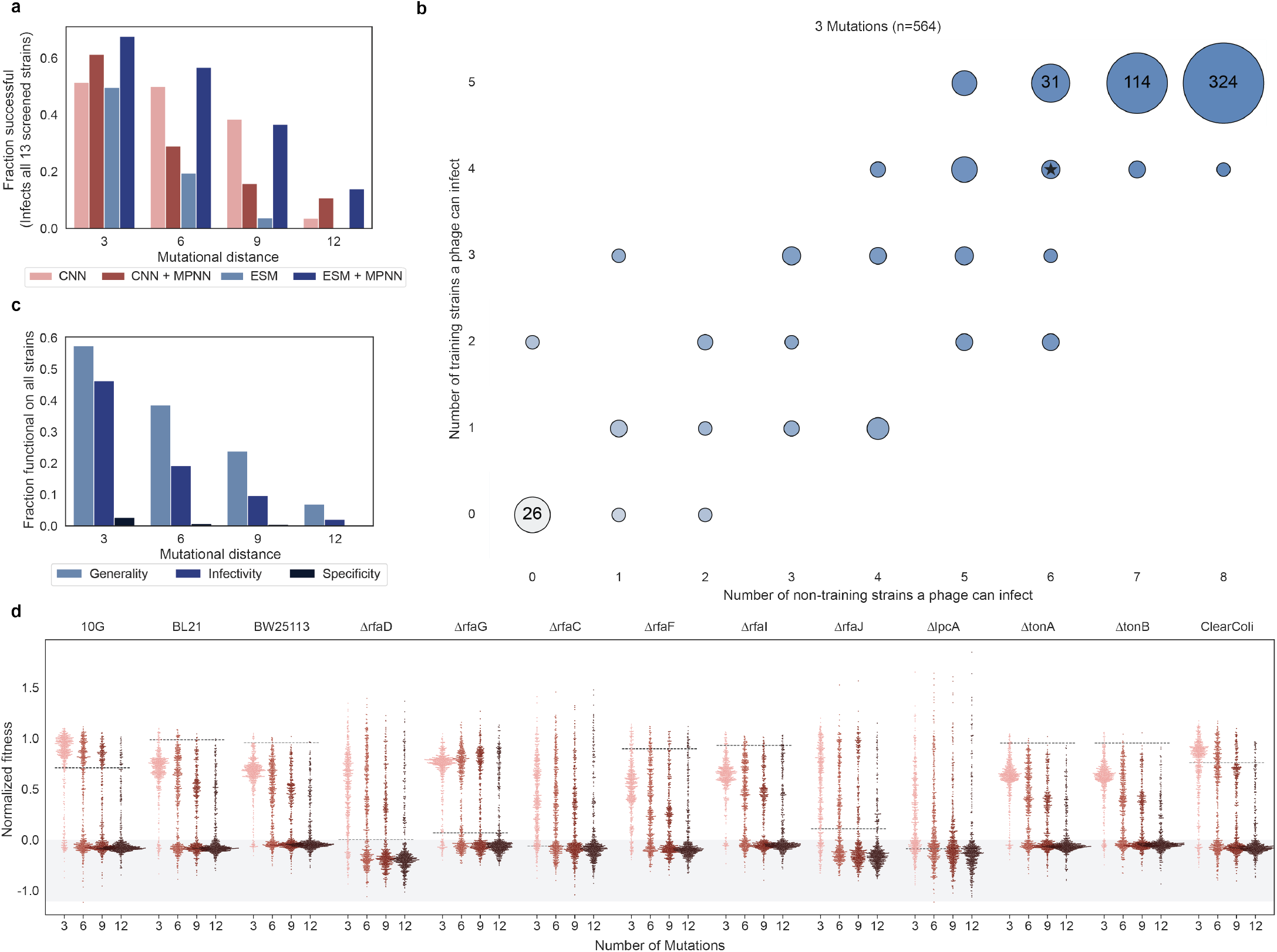
Phages optimized for generality infect diverse training and non-training receptor deletion hosts. a, Fraction of successfully designed generalist phages that could infect all 13 receptor deletion *E. coli* strains that were tested. b, Number of training and non-training receptor deletion strains each phage could infect. The size of the circles indicates the number of observations, ranging from 1 to the maximum. The color gradient also increases with the total number of strains a phage can infect. Wild-type is denoted by a star. c, Fraction of all designed sequences that behave as generalists for each design category. e, Normalized fitness (see methods) of all designed generalists on all training and non-training receptor deletion strains. The black dashed line represents the wild-type score for each host strain. Points in the grey region represent non-functional phages.

The distribution of fitness scores for generalist phages was shifted below wild-type for susceptible hosts and above wild-type for resistant hosts (Fig. 5d). This indicates that, as with specificity, there is a fitness tradeoff for multi-host targeting compared to optimization for single strains. The most common mutation for all generalists was G480K, which occurred in ∼50% of designs and was also identified to be highly beneficial for 3/5 of the host strains in the single mutant dataset^8^ (Supplementary Fig. 10f). Other than this one common mutation, the successful generalist phages were highly diverse, and mutations were not exclusively localized to the primary binding interface of the RBP but rather spread throughout the protein. The diversity of successful mutations throughout the entire sequence suggests that the neural networks extracted non-intuitive mutational trends from the sequence-function data to generate new mutation pairings.

### Secondary validation of infectivity and specificity for designed phage

Due to the nature of selection-based sequencing assays, the fitness scores for a given phage are a function of all library members. This effect, combined with the exponential amplification of active phage, leads to high variability in the scale of scores measured in the high-throughput assay. As a result, the scores do not represent an absolute measurement of virulence, but rather a relative measure constrained by the library and length of incubation. In this work, we conducted optimizations based on these relative fitness scores and observed high success based on the same relative selection-based sequencing assay. To validate the absolute specificity of the designed phage, we selected a handful of designed phages for clonal preparation and screening with a plate-based plaque assay, in which the number of plaques was counted in duplicate on relevant hosts and compared to a reference host to compute the efficiency of plating (EOP).

For several variants, we observe significant improvements in virulence relative to wild-type T7 RBP (Supplementary Fig. 12). For example, a clonal phage for objective xxxx1 has a 1.7×10^4^ fold improvement in EOP compared to wild-type T7 (Supplementary Fig. 12a, p<0.0005). For a nine mutant generalist phage (11111), we observed a 5.0×10^3^ fold increase in the minimum EOP on the five *E. coli* strains that the variant was designed toward (Supplementary Fig. 12f, p<0.05). This improvement was achieved by maintaining virulence on the susceptible 10G, BL21, and BW25113 strains at levels similar to wild-type, while introducing mutations that improved virulence toward the resistant ΔrfaD and ΔrfaG strains 5.0×10^3^ and 110 fold, respectively.

In optimizing for specificity, we selected mutations predicted to maximize the virulence gap between target and anti-target strains. Notably, this strategy permits decreases in virulence on target strains if these effects enable greater depletions in virulence toward the anti-target hosts. This effect is apparent for a six mutant from the objective x101x, which had a 1.7×10^4^ fold improvement in specificity over wild-type (Supplementary Fig. 12c, p<0.0005), but also incurred a ∼five-fold decrease in virulence toward one of the target strains. Similarly, a variant from objective 11011 showed a decrease of 12x toward the 10G target strain and a decrease of 15x toward the BL21 target strain, yet these depletions allowed a striking net 2.7×10^6^ fold increase in specificity relative to T7 (Supplementary Fig. 12e, p<0.05).

Our validation also highlights the constraints imposed by relative fitness measurements in interpreting outcomes from selection-based sequencing assays. For challenging design objectives, such as xx01x and x0011, the selected variants did not display absolute specificity in plaque assays, with EOPs on anti-target strains exceeding those on target strains. Nonetheless, our models produced variants that exhibited substantial relative gains in virulence toward target over anti-target strains relative to wild-type: 9.5×10^4^-fold (Supplementary Fig. 12b, *p*<0.0005) and 590-fold (Supplementary Fig. 12d, *p*<0.005) for xx01x and x0011, respectively. These results reflect the inherent limitations of using relative selection-based data to predict absolute function. Despite this constraint, our models consistently designed variants that either achieved absolute specificity or improved selectivity by several orders of magnitude, validating the effectiveness of leveraging insights from multifunctional sequence data for specificity design.

### Discussion

In this work, we achieve high success rates for the multitask machine learning-directed design of bacteriophages for a diverse set of targeting objectives, even when mutating up to ∼15% of the considered terminal RBP domain relative to wild-type (Fig. 2b). When optimizing for a single task we find that models trained on greater than tens of thousands of fitness measurements do not benefit significantly from additional information in the form of structure or protein language models in terms of the design success rates, unlike that which has been shown for learning from small sequence-function datasets^24,36^. It appears that fine-tuning ESM2 did not significantly impact design success compared to training a simple CNN (Fig. 3a). In a post hoc analysis we compare model predictive performance when all layers of ESM2 are fine-tuned with parameter efficient fine-tuning^37^, which improved classification accuracy relative to our original fine-tuning method (Supplementary Fig. 13). Regardless, reduced or equivalent ESM2 performance both suggest that the next token prediction task with which ESM2 was trained may not be useful for maximizing protein function when fine-tuned on large datasets, which is consistent with other reports that protein function prediction from fine-tuned language models does not scale with model size^38,39^.

While additional information from structure and protein language models did not improve the ability to design single-task infectivity, including these models was useful for complex multiobjective tasks, particularly when one or more targets are minimized (Fig. 4a, Supplementary Fig. 7a,b). This likely arises from the fact that models trained on sequence-fitness data do not inherently recognize the existence of a lower limit for protein function, resulting in continuous *in silico* minimization even when a variant is fully non-functional. By constraining design with structure, protein language models, and/or modified score functions, this over-minimization can be mitigated.

We show that different models appear to make different mistakes (Supplementary Fig. 5a) and that prediction errors were more likely to occur at less frequently mutated positions (Supplementary Fig. 5b). These observations suggest notions that may be useful for future engineering tasks on alternative systems, such as combining multiple modeling approaches to reduce individual inaccuracies of each model and leveraging the frequency that a model incorporates mutations as a filter for design.

Because the models in this paper were trained on a defined set of host targets, they could only make predictions for how RBP mutants would behave toward these five strains. However, we show that this limitation can be overcome by optimizing for generalist function, or high fitness on all known strains, which enabled the identification of improved fitness over wild-type T7 on unseen hosts (Fig. 5c). We see this approach of designing generalists as a useful method to synthesize semi-guided libraries for new tasks without requiring a round of unguided initial design with random or site saturation mutagenesis libraries. Similar approaches would be particularly useful to engineer or diversify enzyme function, for which retention of dynamics and catalytic residues need to be fixed while making subtle changes in substrate positioning and binding specificity^40,41^.

Designing for higher complexity tasks was often much more difficult, especially for objectives that maximized virulence toward resistant hosts while minimizing virulence toward susceptible hosts (Supplementary Fig. 9). This tradeoff was also apparent for the generalist phages, which often behaved worse on susceptible hosts compared to wild-type T7 but demonstrated broader overall promiscuity. While this dataset does not permit broad conclusions about phage evolution, it does provide insight into the local mutational landscape of T7. Our results suggest that minimizing virulence on susceptible hosts may incur additional fitness costs on target hosts, a tradeoff that could be unfavorable in certain evolutionary contexts. In line with previous findings, our analysis of mutation effects across a diverse panel of hosts indicates that the local mutational landscape of wild-type T7 is relatively specialized for the strains we tested^15,16^. The limited host generality observed for wild-type T7 implies that a broad host range is not strongly selected for and that achieving high virulence on at least one host may be more advantageous.

An interesting caveat to the low promiscuity of wild-type T7 is that many designed phages are only a few mutations away from opposite targeting capabilities. This result is consistent with existing hypotheses in the field of phage evolution that host targeting plasticity may have evolved as a mechanism for phages to maintain virulence as their host evolves or when the primary host is no longer proximal^42,43^. This high evolvability also presents a challenge for replication-competent phage therapeutics because individual phage variants exist as quasispecies of a consensus phage, enabling rapid evolution of phage toward unintended hosts *in vivo*. In the future, directed design of phages with higher sequence diversity may be necessary to overcome this constraint and advance bacteriophage toward clinical adoption.

In summary, we demonstrate the application of ML-guided protein engineering to directly design multiple protein functions by optimizing phage infectivity, specificity, and generality. As more data describing variant-host interactions are collected, these methods can be leveraged to design higher complexity specificities or toward new hosts. Directed design of bacteriophage targeting specificity could eventually be used to design bacteriophages for alternative antibiotics, precision microbiome editing, and biosanitation^11,17,44,45^. However, we also hope that this work illuminates the capabilities of machine learning to understand complex multifunctional protein landscapes and navigate across multiple optima, which is of critical importance for protein classes such as enzymes, GPCRs, transporters, antibodies, and transcription factors, among many others. While many protein engineering efforts rely on the collection of low-N datasets, we advocate for the use of high-throughput experimentation to collect data at the scale of >10^4^ perturbations, ideally under multiple pressures or conditions. When paired with machine learning, these large-scale datasets will enable higher design success rates, deep sequence diversification, the design of complex specificities, and overall help advance methods for machine learning-directed design of protein function.

## Methods

### Machine learning model training and implementation

CNNs were implemented in PyTorch and trained from random initializations. The 35 million parameter model from the ESM2 suite was fine-tuned by appending a two-layer fully connected network after the embedding block and allowing training of the final embedding layer and these appended neurons. An ensemble of 100 CNNs and an ensemble of 10 fine-tuned ESM2 models were trained for downstream phage design. Each model in the ensembles was trained with different random initializations. 80-10-10 training, validation, and test splits were used for each model, with the test dataset remaining constant for all dataset splits. A weighted MSE loss function was implemented based on the fitness scores and the inverse of the replicate standard error. This approach biases the model the fit closer to higher-quality data and reduces overfitting. For the thresholded regression, a threshold was set manually for each strain and dataset based on the distribution of fitness scores at the score below which phage was expected to be fully nonfunctional. Weights for scores below this threshold were set to the mean weight for each output to ensure the models did not overfit to functional variant data. Strains and each dataset were organized as separate outputs due to scaling differences across datasets. To determine the final predicted scores for each strain, the relevant scores for each dataset were averaged in a weighted fashion, where weights for each dataset were optimized during hyperparameter optimization. All hyperparameters were optimized using Optuna^46^.

### Processing and generation of training data

This work utilizes three different datasets describing the impact of mutations of phage virulence on a panel of *E. coli* hosts. Two of these datasets were previously published. The first is a site saturation mutagenesis library which contains all single mutations in the terminal 82 residues of the T7 RBP. This phage library was screened against 10G, BL21, BW25113, ΔrfaD, and ΔrfaG. A second dataset was collected as part of this work to augment the amount of training data available (Supplementary Fig. 1). We refer to this library as a combinatorial library, which was screened against 10G, BL21, BW25113, and ΔrfaG. The combinatorial library contains 12,830 combinations of double and triple substitutions of 122 individual substitutions throughout the terminal 82 residues of the T7 RBP. Substitutions were selected based on activity profiles in the original five hosts tested during site saturation mutagenesis of the tip domain. The top 25 candidates enriched on all five strains and a series of 20 substitutions that demonstrated enrichment on between one to four strains and depletion on the remainder of strains were used to generate the list of 122 candidate substitutions. The third library includes mutations spanning the same region of the RBP, but containing up to 10 mutations identified from metagenomic motifs. This library was screened against a large panel of *E. coli* strains, which includes data for 10G, BL21, BW25113, ΔrfaD, and ΔrfaG. To help normalize distribution variance across strains, data for each strain was z-score normalized using SciPy^47^.

### Variant library generation

Variants were generated over 25,000 iterations of simulated annealing based on model predictions and iterative sampling of random mutations. A desired edit distance was specified before the search and a decision to replace an existing mutation was made at each step. Beneficial mutations were always incorporated, and deleterious mutations were accepted with a probability that decreased over the annealing cycle. Median predictions from the CNN and ESM2 models were used for all optimization tasks. The mean ProteinMPNN predicted probability of each residue fitting the wild-type backbone was used to guide optimization for MPNN-constrained models^18^ (Protein_MPNN_v_48_010 weights). When ProteinMPNN was used to guide the optimization of sequences, the product of the CNN or ESM2 infectivity/specificity score and ProteinMPNN scores was used as the objective function. For infectivity objectives, the predicted score for the strain of interest was maximized directly without consideration of the predicted scores for the other strains. For specificity objectives, the gap difference between the lowest score to be maximized and the highest score to be minimized was used to guide mutation sampling*. Generalists were optimized based on the minimum predicted score of all 5 strains on which the model was trained.

*After sampling several different scoring functions for specificity, we selected an optimization function based on a utopia logistic gap function. This function balances the trade-off between maximization and minimization while theoretically preventing over-optimization once a reasonable score is achieved in either direction, using a logistic (sigmoid) transformation. Before each operation, the sigmoid transformation is applied to normalize each score, ensuring that the true maximum and minimum scores are 1 and 0, respectively. Then, the following operation is used to compute the specificity score.

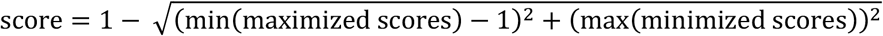

### Phage library preparation

Genes encoding the mutant 82 amino acid terminal region of the RBP were ordered as an oligo pool from Twist and amplified using PCR. The resulting DNA fragments were cloned into a bacterial plasmid (pHT7Rec1 Template) containing a type IIS landing pad for the gene insert. The resulting library was used in the ORACLE phage library preparation method^8^. The first step of ORACLE is optimized recombination, in which the library is recombined into an acceptor phage with lox71 landing sites. Next, in the accumulation step, the resulting phage library is enriched for phages that have undergone this recombination step by allowing infection of the phage into a strain that contains Cas9 and a guide targeting a site in the genome of non-recombined phage (SC101_pCas9_Target3). During these two steps, the bacterial host expresses the wild-type T7 RBP at high levels to reduce library bias (pHT7Helper1 Pbr mCherry+gp17). The inclusion of this helper plasmid yields a genetically modified phage that presents the wild-type RBP. Finally, the enriched library is expressed by having the accumulated phage library infect a bacterial strain lacking this wild-type RBP helper plasmid for a single replication cycle.

### Microbes and culture conditions

All culturing experiments were conducted at 37°C with shaking in standard LB media. The *E. coli* bacterial strains used in this study are as follows: 10G, BL21, BW25113, BW25113*ΔrfaD*, BW25113*ΔrfaG*, BW25113*ΔrfaC*, BW25113*ΔrfaF*, BW25113*ΔrfaI*, BW25113*ΔrfaJ*, BW25113*ΔlpcA*, BW25113*ΔtonA*, BW25113*ΔtonB*, ClearColi BL21(DE3). Susceptible strains included *E. coli* BL21, a B strain derivative, BW25113, a K-12 derivative, and 10G, a DH10B derivative. All receptor deletion strains are resistant to phage infection and contain truncations at different points in the lipid polysaccharide receptor^48–50^. These deletions were made in the BW25113 strain and include BW25113*ΔrfaD*, BW25113*ΔrfaG*, BW25113*ΔrfaC*, BW25113*ΔrfaF*, BW25113*ΔrfaI*, BW25113*ΔrfaJ*, BW25113*ΔlpcA*. Deletion of these genes can reduce the ability of T7 to infect *E. coli* by several orders of magnitude^16^. ClearColi BL21(DE3) contains a full deletion of the lipid polysaccharide in the BL21 background strain. BW25113*ΔtonA* and BW25113*ΔtonB* represent deletions in genes encoding an iron transporter, which serves as a secondary receptor for T7. Deletion strains are referred to throughout simply as the gene that was deleted for brevity. No antibiotics were used during the selection experiments, but during library preparation, Kanamycin50 or Kanamycin50+Spectinomycin115 were used.

### Growth-based selection of phage

Bacterial cultures were grown to an OD600 of ∼0.4 before adding the phage library at a multiplicity of infection (MOI) of 0.01 and shaking at 37°C for two hours (or until culture lysis for ΔrfaD, ΔrfaG, ΔlpcA). After this selection, the cultures were placed on ice to halt further infections, and a small amount of chloroform was added to the culture to kill any remaining bacteria. Post-selection phage populations were harvested by filtering the lysate through a 0.22µm filter.

### Next-generation sequencing and data processing

Post-selection phage populations were used as a template for a PCR (KAPA HiFi, KK2102) directly off the phage genome using primers that targeted the mutated region in the RBP. These primers also added indexes corresponding to the biological replicate. A second PCR was conducted using the first PCR as a template which added i5 and i7 indexes for Illumina sequencing. Sequencing was done at the UW-Madison Biotechnology Center on a NovaSeq X Plus shared 2×150 run. The resulting data was processed using custom Python scripts to extract the sequences of input and count the number of observations for each designed variant. Once pre and post-selection counts for each variant were available, variants were filtered to exclude sequences with less than 25 pre-selection counts or less than 5 post-counts for all of the strains screened. Then, scores were calculated using the log ratio of post and pre-selection counts and maximum likelihood estimation to combine biological triplicates^51^. All analyses utilize or show the combined triplicate scores. Count correlations for nearly all replicates frequently exceeded Pearson correlations of >0.95, except in instances of post-selection measurements where variant dropout was frequent. For data shown in this study, scores were z-score normalized^47^ and additionally normalized between zero and one using the manually set threshold for phage function and the 98^th^ percentile score for that strain. This normalization approach was chosen so that zero would approximately represent non-functional phage and 1 would represent the maximum observed fitness on that strain (excluding outliers).

### Computational analysis and statistical tests

Plotting and analysis were done using seaborn and matplotlib^52,53^. Weighted Spearman metrics were calculated using a Python implementation of the weighted Spearman R package where the weights were the same weights used during training^54^. Dimension reduction was done on one-hot encoded RBP variants using PaCMAP^55^ with the default parameters, except for n_neighbors and MN_ratio which were set as 100 and 2, respectively. The coordinates of the dimension reduction were saved and reused in all dimension reduction figures. In Fig. 1c and 1d, circled regions were placed during interactive plotting, and the sequences were determined based on the coordinates of points within the circle. These circled regions were selected for their high concentration of sequences designed by each model and high abundance of generalist designs (to reduce motif divergences caused by differences in objective). A Mann–Whitney U test statistical test was used to compare distributions of BLOSUM scores. A Fisher’s Exact Test was used to compare the success summed across sub-objectives for all pairs of models and mutational distances for infectivity, specificity, and generalist tasks. A student’s t-test was used to determine the significance between conditions in clonally validated phage data. During some model comparisons, we use the average precision (AP) metric to compare classification accuracy due to unbalanced datasets. This metric is calculated by taking the area under the precision-recall curve, where precision is defined as (true positives)/(true positives + false positives), and recall is defined as (true positives)/(true positives + false negatives)^26^.

### Plaque assays

Plaque assays were performed to estimate phage titers by mixing phage within a top agar solution of 0.5% agar with 250µL of overnight 10G+Helper bacterial culture. Phage was added at a dilution to enable plaque counting. These plates were incubated for 16 hours before counting plaques. The titers determined through plaque assays were used to guide the amount of phage required to achieve target MOI during selection experiments and for clonal phage validation of experimental results.

### Construction of clonal phage variants

To make clonal phage variants for validation, Gibson Assembly was performed using the NEBuilder HiFi DNA Assembly Master Mix (NEB E2621L). The T7 phage genome was split into five ∼10kb fragments and mixed at a 1:1 molar ratio along with a synthetic gBlock carrying the gp17 variant sequence. Following the Gibson Assembly, drop dialysis was performed using a 0.025µM mixed cellulose ester membrane (VSWP01300) and *E. coli* 10G cells were then transformed using electroporation. The transformation mix was then recovered for 1 hour at 37°C before 200µl of the transformation mix was directly plated in 0.5% LB+10G top agar. Following overnight incubation at 37°C, individual plaques were picked and sequenced. Codon usage for mutations was determined by selecting codons optimized based on the T7 genome. Mutations used for clonal validation are noted in the table below. Only the terminal 82 residues of the RBP were mutated. Mutations are numbered according to the position in the full T7 RBP.

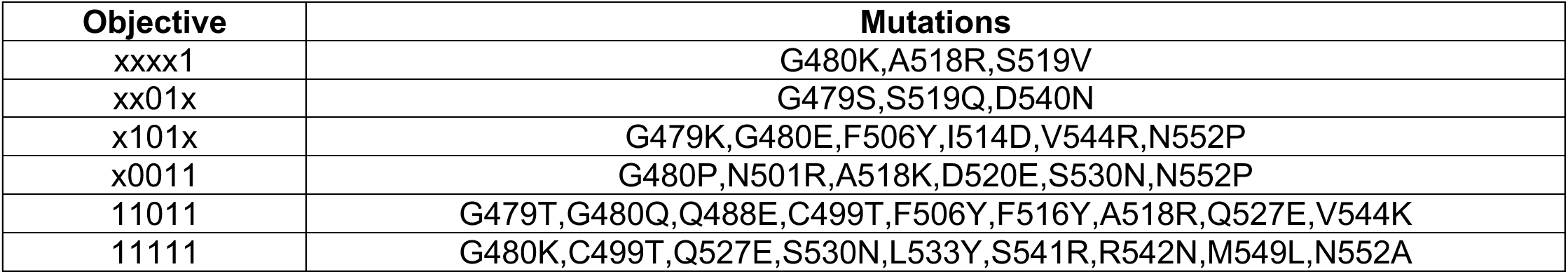

| Objective | Mutations |
| --- | --- |
| xxxx1 | G480K,A518R,S519V |
| xx01x | G479S,S519Q,D540N |
| x101x | G479K,G480E,F506Y,I514D,V544R,N552P |
| x0011 | G480P,N501R,A518K,D520E,S530N,N552P |
| 11011 | G479T,G480Q,Q488E,C499T,F506Y,F516Y,A518R,Q527E,V544K |
| 11111 | G480K,C499T,Q527E,S530N,L533Y,S541R,R542N,M549L,N552A |

## Author Contributions

NN, PH, and SR conceptualized and designed the study. NN, PH, and SE performed wet lab experiments. NN and PR developed and implemented the machine learning models and designed computational analyses. NN drafted the manuscript, and all authors reviewed and approved the final version.

## Acknowledgments

This work was supported by the NIGMS Biotechnology Training Program T32GM135066 (N.N.), NSF CAREER 2237251 (S.R.), and Defense Threat Reduction Agency HDTRA12310023. The authors thank Sam Gelman for valuable discussions and for providing code related to machine learning methods and fine-tuning. We also acknowledge the UW Biotechnology Center for providing next-generation sequencing services.

## Data and code availability

Data and code used in this work will be made available upon publication. Code for training machine learning models on protein sequence-function data is formatted as a flexible framework as a tool for the research community.

