## Supplemental figures for "Multiobjective learning and design of bacteriophage specificity"

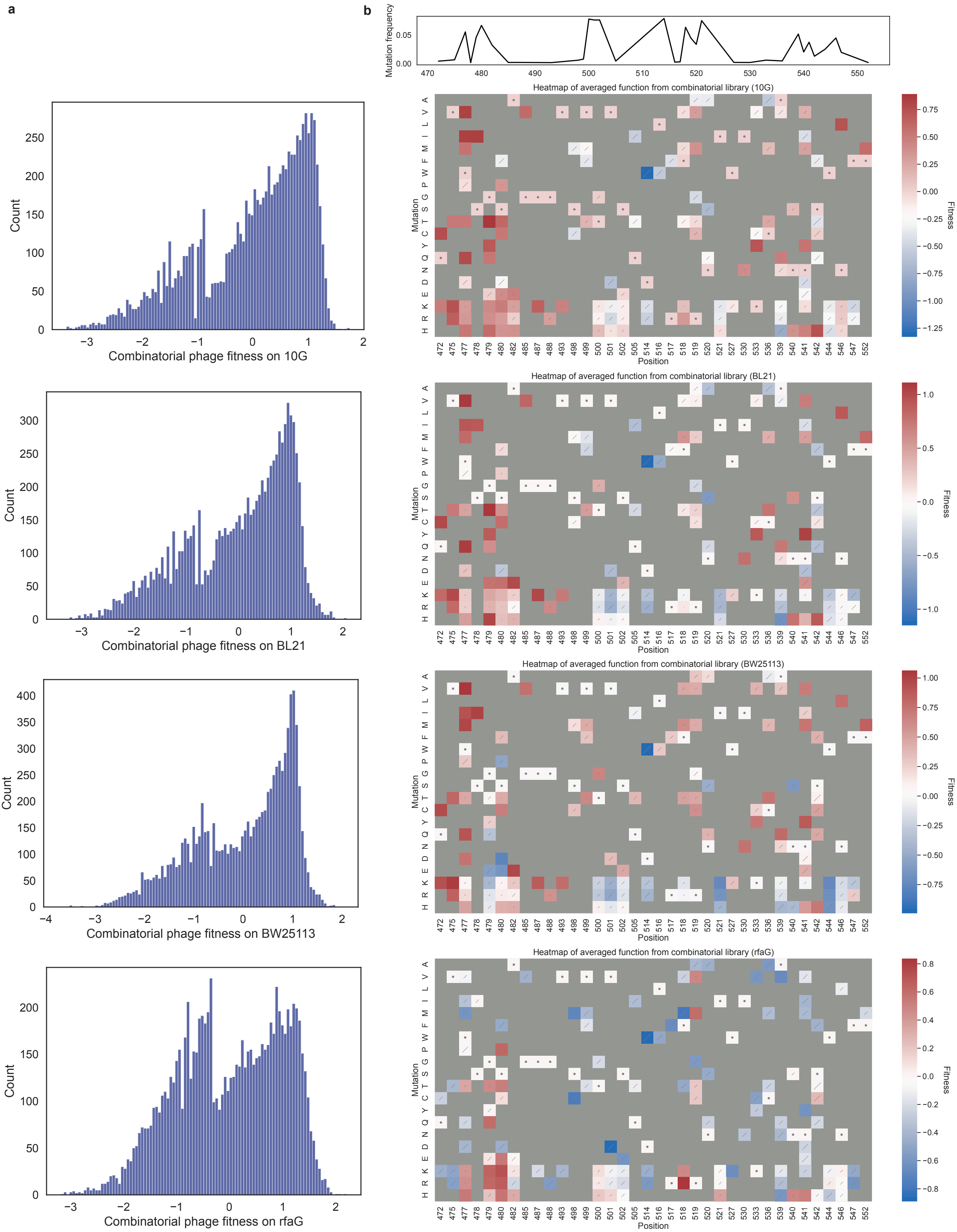


**Supplementary Fig. 1: Fitness profile of combinatorial phage library**

a, Histograms of fitness measurments for combinatorial phage variants on 10G, BL21, BW25113, and ΔrfaG. The combinatorial library was not screened against ΔrfaD. b, DMS heatmaps of averaged fitness effects from the combinatorial library on each strain. WT is indicated by a dot and the standard error between tripllicates is shown with a bar. Grey boxes were not included in the library. Mutation frequency at each position is shown above the heatmaps.

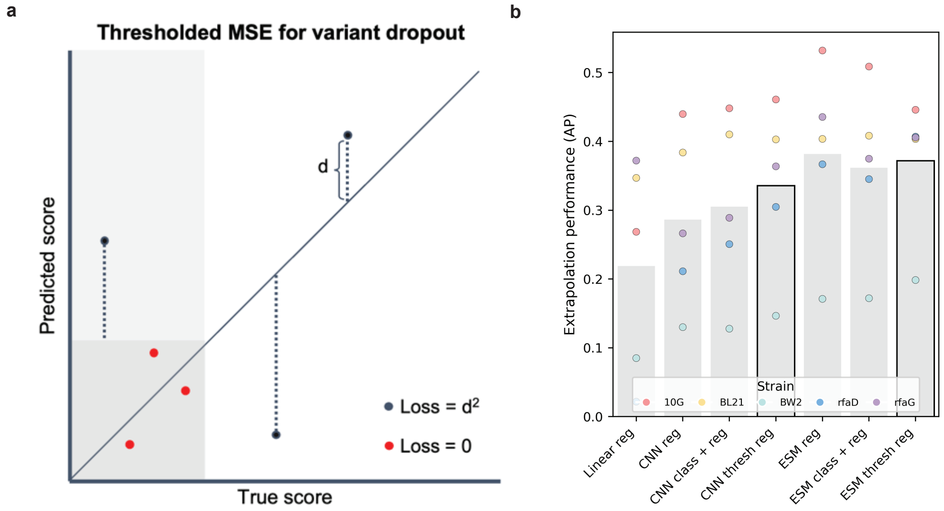


**Supplementary Fig. 2: Additional model guided-design details**

a, Schematic of the custom thresholded mean square error loss function used in this study. Thresholds for the minimum detectable functional score are manually set for each strain and dataset. b, Several different architectures and learning methods were sampled to evaluate the performance of different approaches when models were trained on 1-3 mutant data and tested on higher order mutants. The CNN and ESM thresholded regression were used for downstream sequence design based on these results.

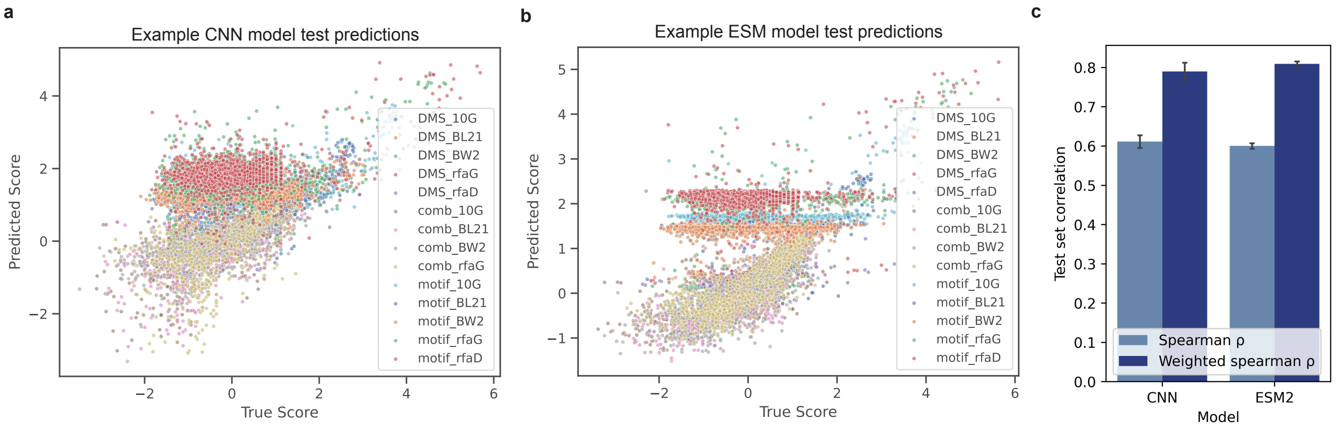


**Supplementary Fig. 3: Model performance on test set**

Scatterplots of predictions from CNN (a) and fine-tuned ESM (b) models on held out test dataset. c, Performance of CNN and ESM models used for sequence design on held out test sets. Weights for spearman correlation are derived from the inverse of the standard error determined from maximum likelihood experimental replicate combination.

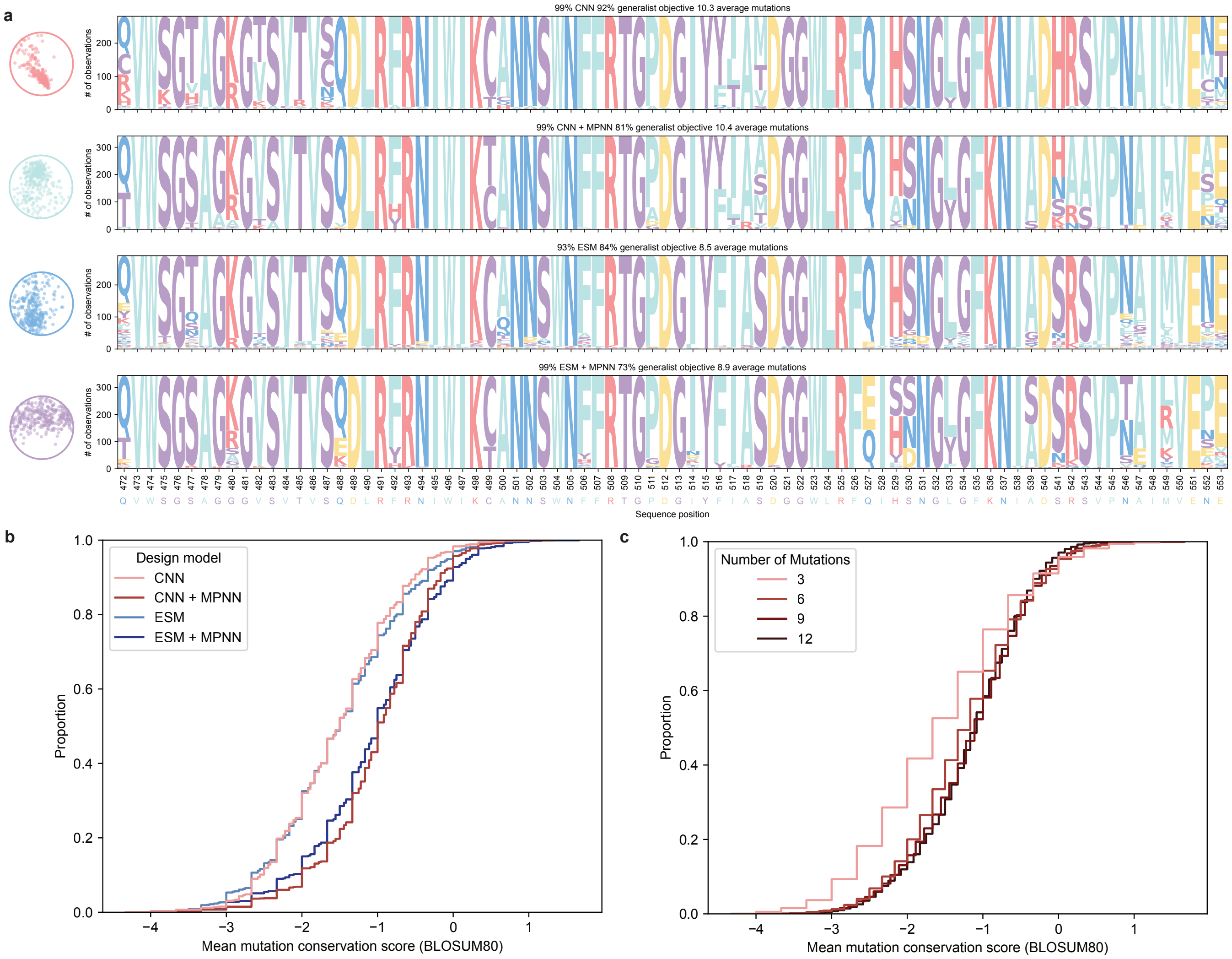


**Supplementary Fig. 4: Choice of design model influences mutation preferences and conservation**

a, Full motif diagram of sequences circled Figure 1c. b, Mean BLOSUM80 for mutations by model (higher is more conserved). c, Mean BLOSUM80 of mutations by mutational distance.

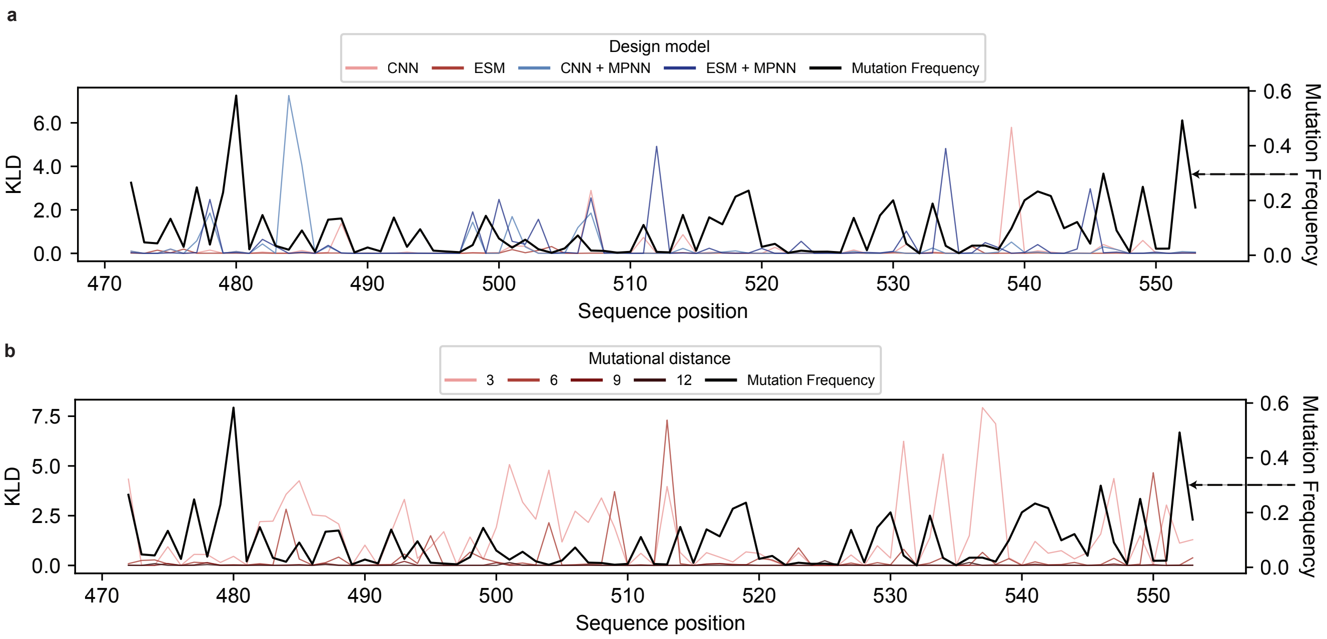


**Supplementary Fig. 5: Positional design error by model and mutational distance**
Positional Kullback-Leibler Divergence (KLD) between mutation frequency distributions for non-functional variants compared to the distribution of all variants as the reference for design model (a) and mutational distance (b). Right axes show the overall mutation frequency at each position.

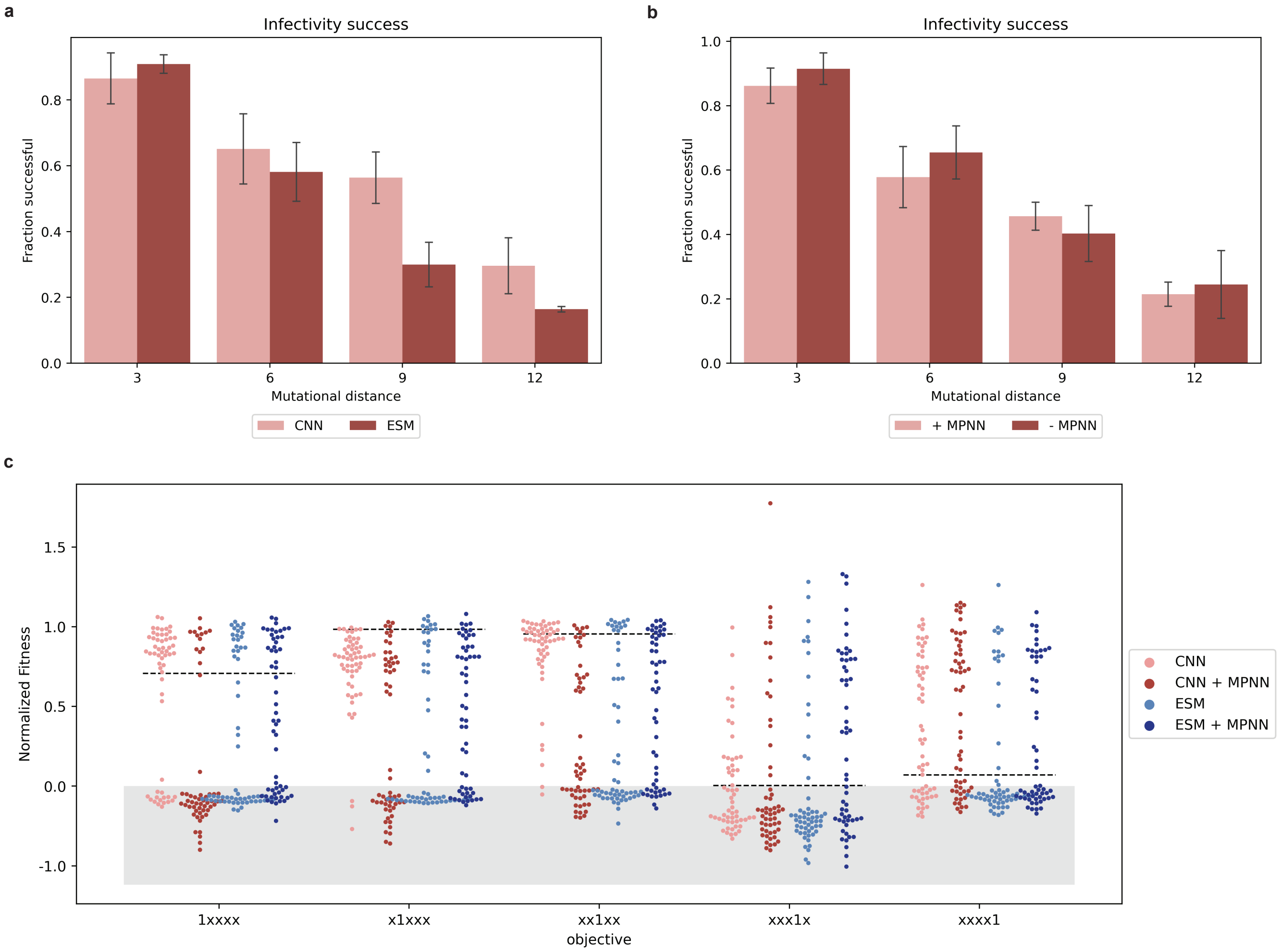


**Supplementary Fig. 6: Additional characterization of phage infectivity**

a, Infectivity design success with respect to the base fitness model used. b, Infectivity design success considering whether or not structural constraints from ProteinMPNN were added during sequence design. c, Normalized fitness for all phages designed for the infectivity category. Plot hue indicates the model used to design phages. All mutational distances are included in this analysis.

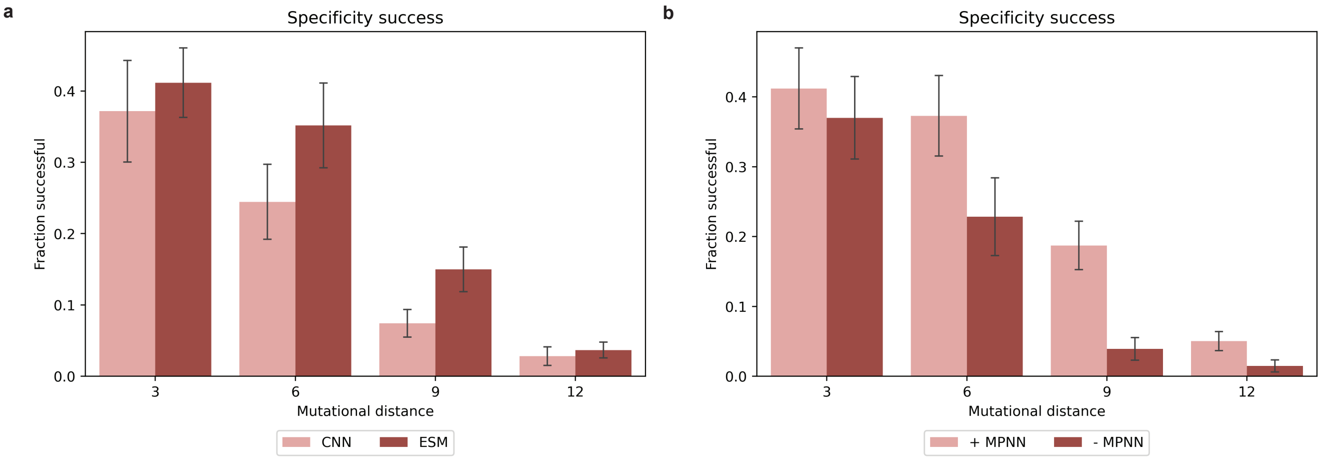


**Supplementary Fig. 7: Additional characterization of phage specificity**
a, Specificity design success with respect to the base fitness model used. b, Specificity design succes considering whether or not structural constraints from ProteinMPNN were added during sequence design.

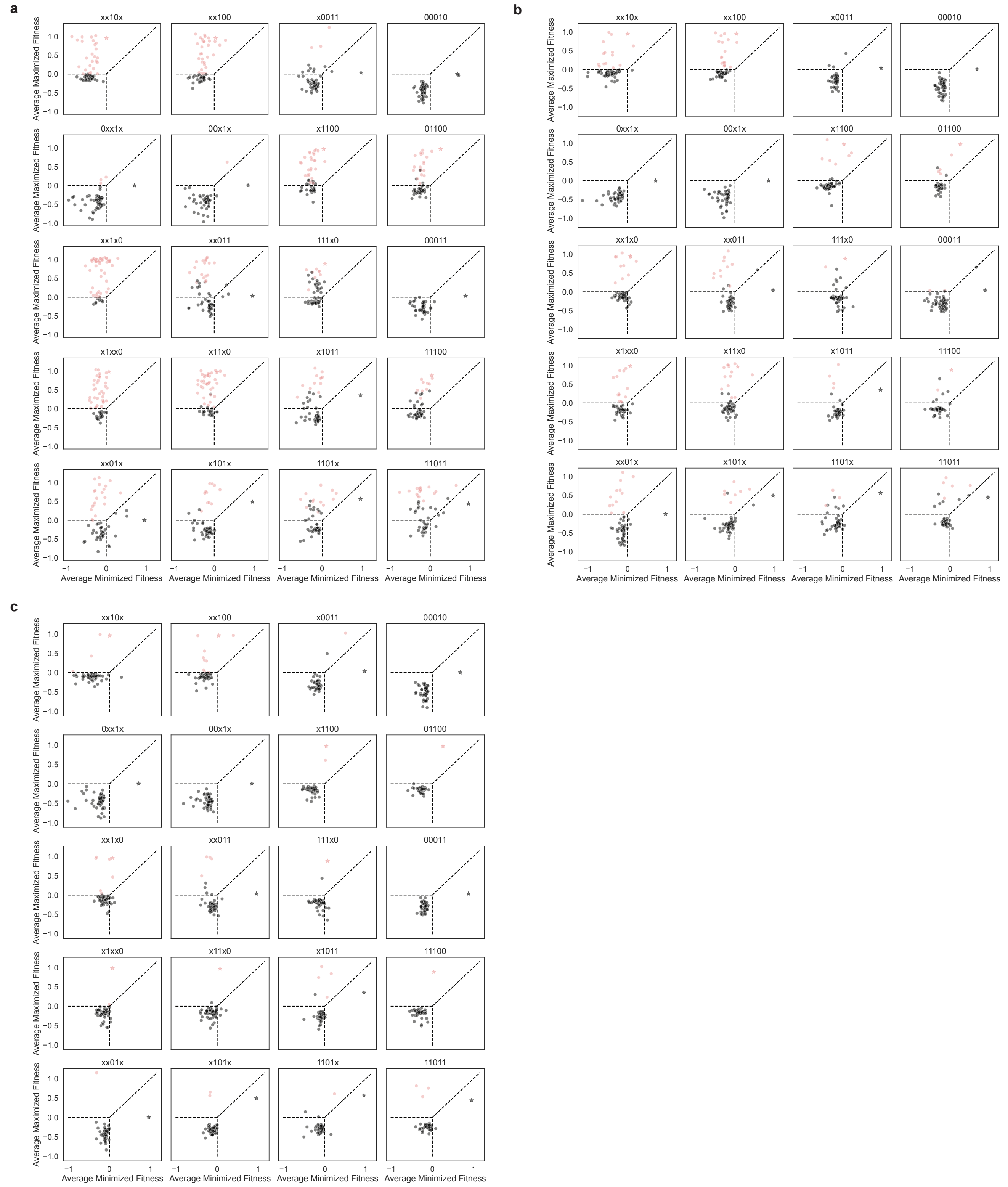


**Supplementary Fig. 8: Analysis of designed phage specificity at higher mutational distances**

Average fitness of mutant phages on the maximized and minimized strains for each specificty objective for 6 (a), 9 (b), and 12 (c) mutations from WT. Red points meet all criteria for a particular specificty and black points were unsuccessful for one or more condition. WT fitness is indiced by a star.

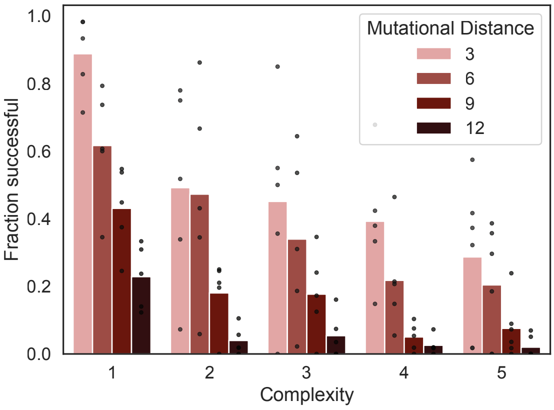


**Supplementary Fig. 9: Impact of complexity on design success**
Success rates by the complexity of the design objective (how many targets were considered for either maximization or minimization during optimization). Average of the success for all objectives at the indicated compexity and mutational distance. Success rates for all objectives are overlayed as black points.

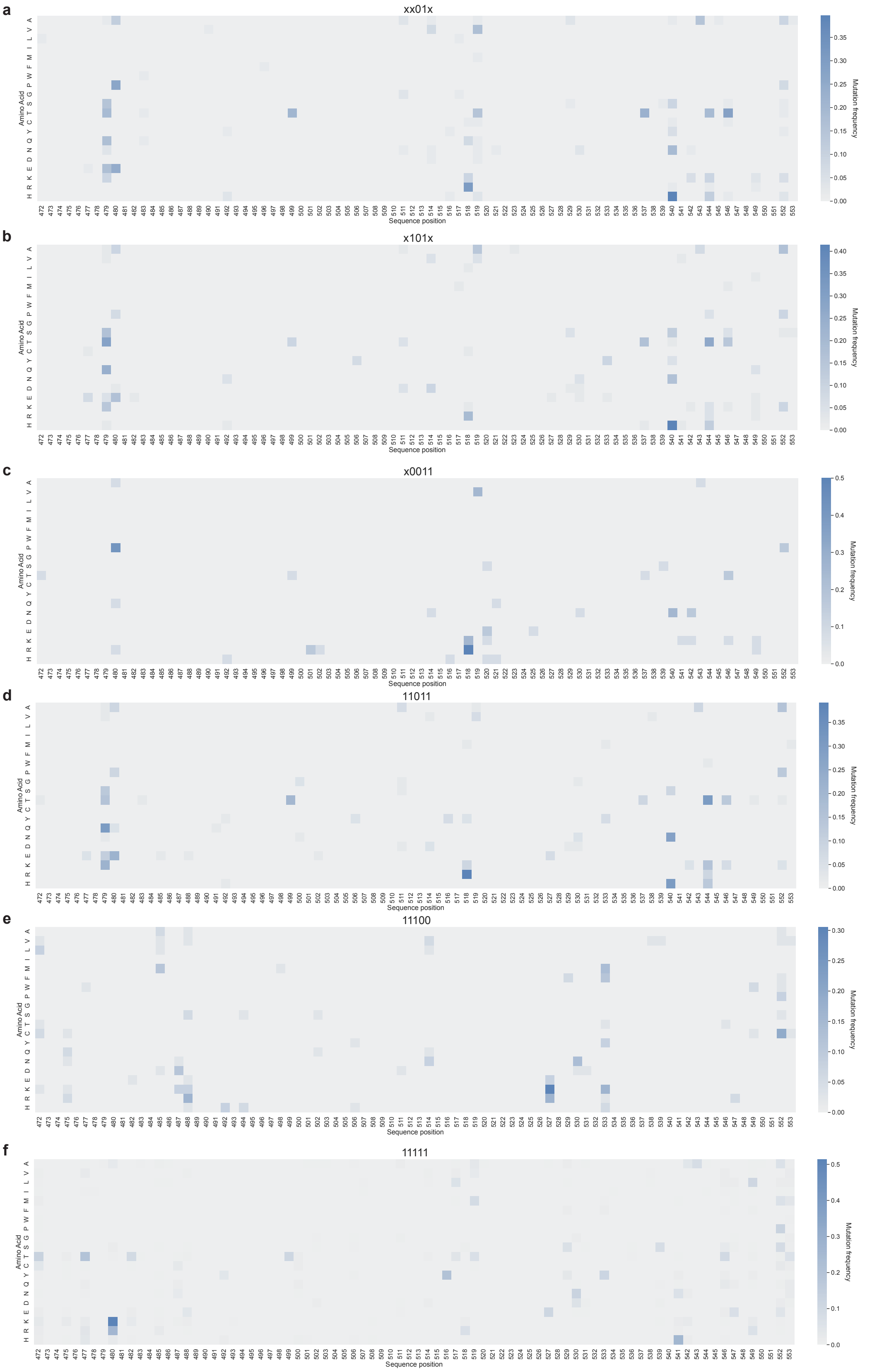


**Supplementary Fig. 10: Mutation frequency heatmaps for selected objectives**
Heatmaps of mutation frequencies for amino acids at each position. WT frequencies are excluded from the calculation and frequencies represent the fraction of phage for each objective that were successful and contained that particular mutation.

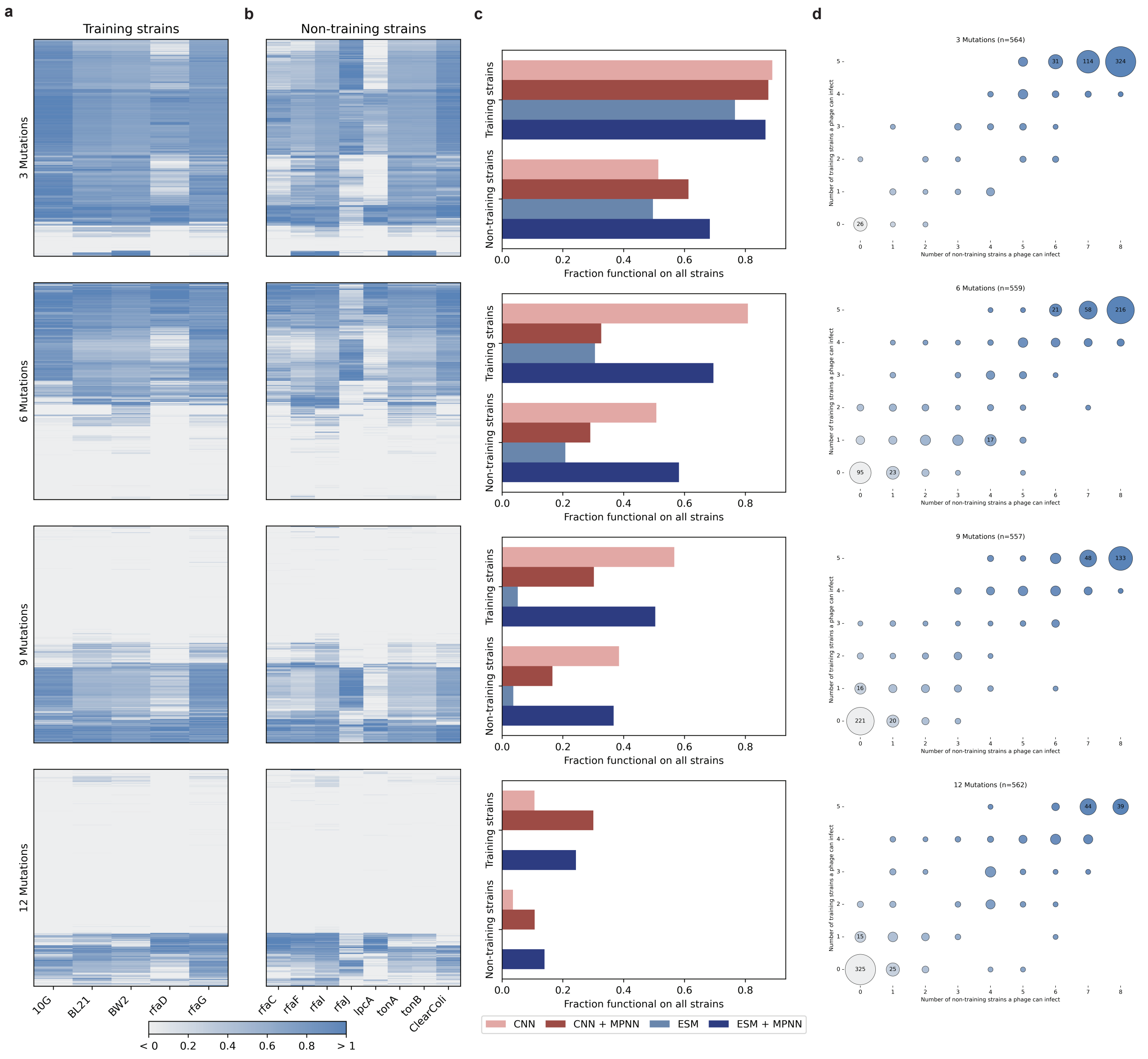


**Supplementary Fig. 11: Additional characterization of phage generality**

a, Clustermaps of fitness scores for designed phages on training (a) and non-training (b) receptor deletion strains. Each row in the clustermaps represents a single phage. c, Fraction of phages that could infect either all training strains or all non-training receptor deletion strains. Plots are ordered vertically by mutational distance relative to WT. d, Number of training and non training receptor deletion strains each phage could. Size of circles indicates number of observations, ranging from 1 to the maximum. Color gradient increases with the total number of strains a phage can infect.

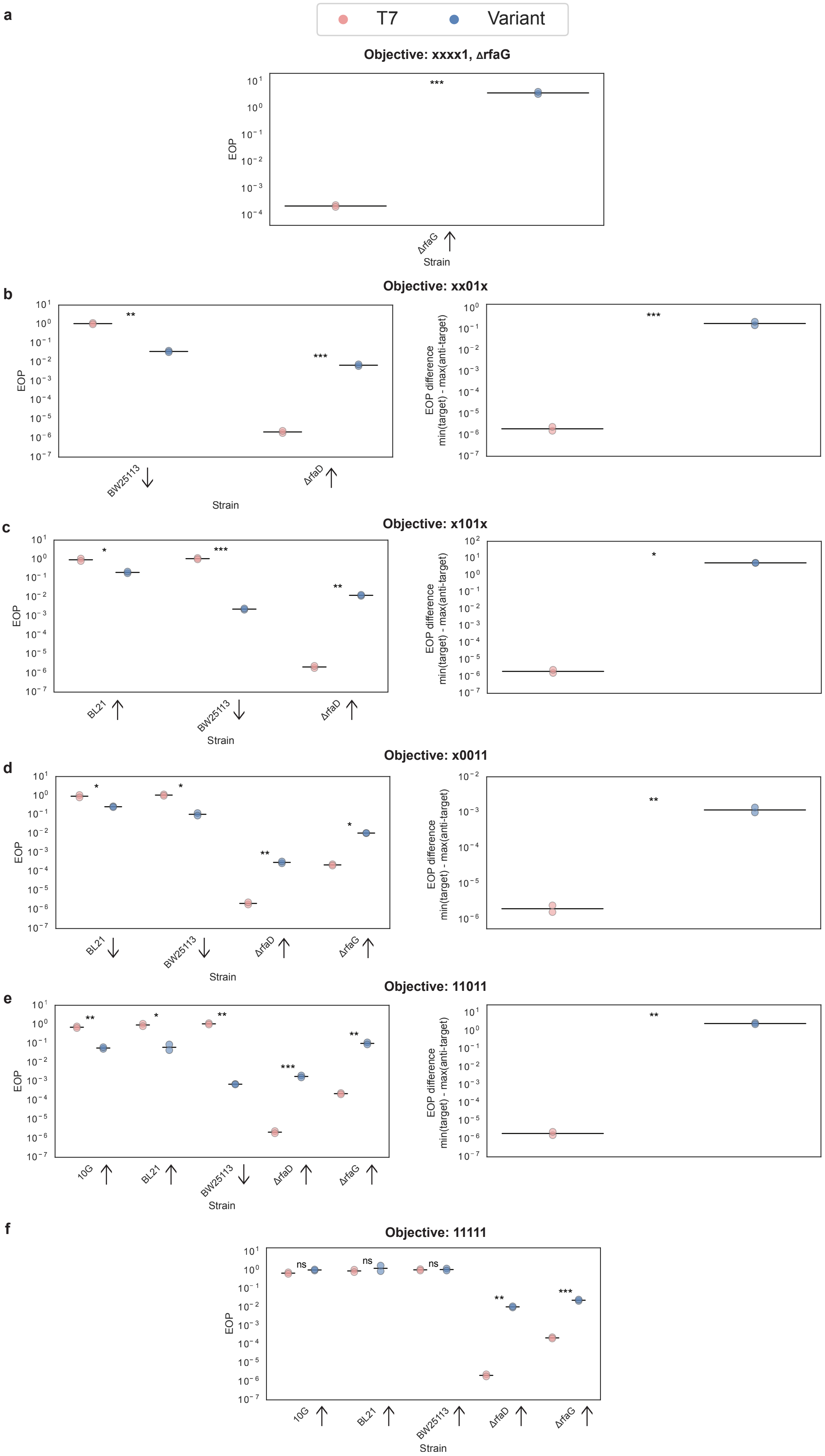


**Supplementary Fig. 12: Validation of engineered phage in clonal experiments**
Plaque assays were performed in duplicate on each relevant host for a variant phage of select objectives and normalized to wildtype T7 phage against the same hosts, where an EOP of zero represents virulence equivilent to T7 infecting 10G+helper (constitutive wildtype T7 RBP expression). Refer to Fig. 1b for legend of specificity codes. Validated objectives include xxxx1 (a), xx01x (b), x101x (c), x1100 (d), 11100 (e), 11011 (f), and 11111 (g). For specificity objectives, the right plot shows the gap difference in observed EOPs for the target objective. A Students t-test was used to determine if the infectivity of the variant phage was significantly different than the WT phage. * indicates p<0.05, ** indicates p<0.005, and *** indicates p<0.0005.

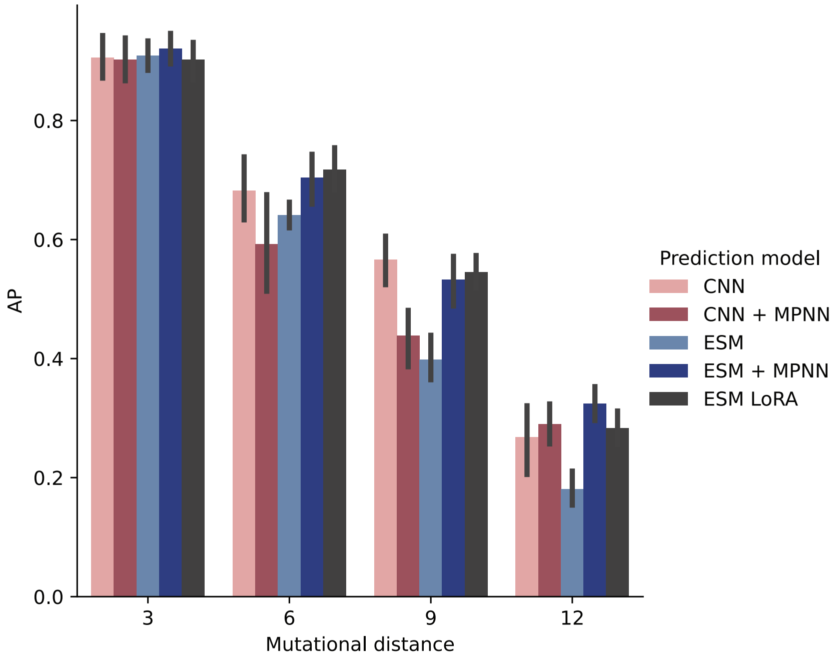


**Supplementary Fig. 13: Comparison of model acuracies to an alternative fine-tuning approach**
Comparison of the area under the precision-recall curve (AUPRC/AP) when each model is tasked with predicting whether all designed sequences meet their intended objective (Same task as in Fig. 2b). ESM LoRA represents the same ESM model fine-tuned using low rand adaptation (LoRA).
